# Identification and characterization of genes encoding the nuclear envelope LINC complex in the monocot species *Zea mays*

**DOI:** 10.1101/343939

**Authors:** Hardeep K. Gumber, Joseph F. McKenna, Amado L. Estrada, Andrea F. Tolmie, Katja Graumann, Hank W. Bass

**Affiliations:** Department of Biological Science, Florida State University, Tallahassee, FL, USA, 32306-4295; Department of Biological and Medical Sciences, Faculty of Health and Life Sciences, Oxford Brookes University, Oxford, UK, OX30BP

**Keywords:** KASH, LINC, Maize, nuclear envelope, SUN

## Abstract

The LINC (Linker of Nucleoskeleton to Cytoskeleton) complex is an essential multi-protein structure spanning the nuclear envelope. It connects the cytoplasm to the nucleoplasm, functions to maintain nuclear shape and architecture, and regulates chromosome dynamics during cell division. Knowledge of LINC complex composition and function in the plant kingdom is primarily limited to Arabidopsis, but critically missing from the evolutionarily distant monocots which include grasses, the most important agronomic crops worldwide. To fill this knowledge gap, we identified and characterized 22 maize genes, including a new grass-specific KASH gene family. Using bioinformatic, biochemical, and cell biological approaches, we provide evidence that representative KASH candidates localize to the nuclear periphery and interact with ZmSUN2 *in vivo*. FRAP experiments using domain-deletion constructs verified that this SUN-KASH interaction was dependent on the SUN but not the coiled-coil domain of ZmSUN2. A summary working model is proposed for the entire maize LINC complex encoded by conserved and divergent gene families. These findings expand our knowledge of the plant nuclear envelope in a model grass species, with implications for both basic and applied cellular research.

**SUMMARY STATEMENT:** Genes encoding maize candidates for the core LINC and associated complex proteins have been comprehensively identified with functional validation by one or more assays for several of the KASH genes.

## INTRODUCTION

In eukaryotic cells, the nuclear envelope (NE) is a structural hallmark that encapsulates the biparentally-inherited genetic material, the chromosomes. The NE is comprised of inner and outer nuclear membranes, with embedded nuclear pore complexes (NPC) and a variety of nuclear envelope transmembrane proteins (NETs). Two NETs, Sad1/UNC-84 (SUN), and Klarsicht/ANC-1/Syne-1 homology (KASH) domain proteins, interact in the perinuclear space to form the core of an evolutionarily conserved Linker of Nucleoskeleton and Cytoskeleton (LINC) complex which spans the NE (Crisp et al., 2006). LINC complexes carry out many functions ranging from organizing the shape and position of the nucleus as a mobile and pliable organelle to direct mechanotransduction to effects on chromatin structure and dynamics in somatic and meiotic cells (Kim et al., 2015; Kracklauer et al., 2013; Meier, 2016; Meier et al., 2017; Razafsky and Hodzic, 2015; Starr and Fridolfsson, 2010; Tamura et al., 2015; Tapley and Starr, 2013). The biological importance of NE functions comes primarily from studies of opisthokonts, and in humans is reflected through the wide range of NE-defective developmental disorders such as laminopathies or envelopathies (Burke and Stewart, 2014; Fridkin et al., 2009; Janin et al., 2017; Razafsky and Hodzic, 2015).

The core components of the LINC complex are the inner nuclear membrane (INM) SUN-domain proteins (Hagan and Yanagida, 1995; Malone et al., 1999) and the outer nuclear membrane (ONM) KASH-domain proteins (Starr and Han, 2002). The SUN-domain proteins are known to interact with chromatin, nuclear lamins, or lamin-like proteins (Haque et al., 2006; Hodzic et al., 2004; Zhou et al., 2015b). The KASH-domain proteins are known to interact directly or indirectly with cytoskeletal structures including microfilaments, microtubules, intermediate filaments, microtubule organizing centers, and organelles (Luxton and Starr, 2014; Starr and Fridolfsson, 2010). LINC complexes are highly conserved in all eukaryotes, yet many of the components have evolved rapidly or recently, limiting the power of sequence conservation to identify functional homologs.

The body of knowledge of the LINC complex comes primarily from studies in opisthokonts, especially metazoans and yeast. However, there exists a severe gap in our knowledge of plant LINC functions, limiting our ability to understand, predict, and modify plants to meet global challenges in agriculture and biorenewable resources (Godfray et al., 2010). The first SUN domain protein recognized in plants, OsSad1, was found as part of nuclear proteomic study in rice (Moriguchi et al., 2005). Its relationship to the *Saccharomyces pombe* Sad1 was confirmed by localization of OsSad1-GFP fusion protein to the nuclear periphery in onion epidermal cells (Moriguchi et al., 2005). Subsequently, two classes of SUN domain proteins were found to be present in plants; first, the CCSD/C-terminal SUN group, in which the SUN domain is near the C-terminus and second, the PM3/Mid-SUN group, in which the SUN domain is centrally located (Graumann and Evans, 2010; Graumann et al., 2014; Murphy et al., 2010; Oda and Fukuda, 2011). More recently, a large number of SUN domain proteins have been identified in many plant species by homology searches (reviewed by Meier, 2016; Meier et al., 2017; Poulet et al., 2017a).

The C-terminal SUN proteins are similar to those in animals, but the mid-SUN proteins are less well studied as a group despite their known occurrence in plants (Poulet et al., 2017a), yeast (Friederichs et al., 2011), Dictyostelium (Shimada et al., 2010) and mice (Sohaskey et al., 2010). Among the conserved roles that SUN proteins play in plants are those involving the maintenance of the shape and size of the nucleus (Graumann et al., 2010; Graumann et al., 2014), plant growth (Graumann et al., 2014), and meiotic chromosome behavior (Murphy et al., 2014; Varas et al., 2015).

The first plant KASH proteins to be described were the WPP-domain-interacting Other components of the plant LINC complex have been described primarily in a plant model system, Arabidopsis, a eudicot species that diverged from the economically important monocot species ~200 MYA (Meier, 2016; Meier et al., 2017; Poulet et al., 2017a; Tiang et al., 2012; Zeng et al., 2014; Zhou et al., 2015a). proteins (WIPs). The WIPs are plant-specific, outer nuclear envelope associated proteins originally discovered as RanGAP-NE anchoring proteins (Xu et al., 2007) and later shown to bind to SUN and possess KASH-like features (Meier et al., 2010; Zhou et al., 2012). A related group of ONM LINC-associated plant proteins are the WPP-interacting tail-anchored (WIT) proteins (Zhao et al,. 2008), which bind directly to WIPs, indirectly to SUN, and also have WPP-interacting domains. WITs possess the LINC-like activity of connecting the NE to the actin cytoskeleton through their association with plant specific Myosin XI-i (Tamura et al., 2013). In addition to the WIPs and WITs, two other categories of ONM KASH proteins have been described in plants, the SUN-Interacting Nuclear Envelope proteins (SINEs) and the TIR-KASH (TIKs), both of which are NETs (Graumann et al., 2014; Zhou et al., 2014).

There are currently three recognized plant-specific categories of genes encoding proteins in the INM or at the intranuclear periphery that interact with SUN or each other to associate with the LINC complex. These encode the INM Nuclear Envelope Associated Proteins (NEAPs) and nucleoplasmic proteins, including Nuclear Matrix Constituent Proteins (NMCPs, also called CRWNs) and the AtCRWN-binding nucleoplasmic protein AtKAKU4s (Ciska et al., 2013; Dittmer and Richards, 2008; Goto et al., 2014; Masuda et al., 1997; Pawar et al., 2016; Wang et al., 2013). While NEAPs and CRWNs can be directly associated with LINC complexes by binding to C-terminal SUN proteins, KAKU4 is indirectly associated with LINC complexes via interaction with CRWNs (Goto et al., 2014). These interconnected proteins are thought to have roles in chromatin architecture and nuclear morphology, placing the LINC complex squarely in genetic and genomic response pathways in plant cells (Poulet et al., 2017b).

There exist profound biological differences and vast evolutionary distances between monocots (grasses) and eudicots. Given this fact, together with the importance of the nucleus and the LINC complex in basic developmental processes, we set out to define and characterize for the first time the entire LINC complex in the crop grass model species, maize (reviewed by Nannas and Dawe, 2015). We include comparisons across phyla ranging from single cellular chlorophyte to advanced land plants, to highlight both conserved and divergent maize LINC complex components.

## RESULTS

### Identification of maize genes encoding LINC or LINC-associated components

We developed a working model, shown in Figure 1, for the maize LINC complex and associated proteins as identified through bioinformatic or experimental evidence. The proteins are depicted as residing in one of the four cellular locations, the cytoplasm, the ONM of the NE, the INM of the NE, or the nucleoplasm. Despite the considerable difficulty in assigning certain orthology for all of the maize LINC components, we have established the following nomenclature for these candidate maize LINC genes. The genes encoding KASH-like and associated ONM proteins are named with the common prefix of *MLK* for *<u>M</u>aize <u>L</u>INC <u>K</u>ASH* followed by a one letter designation to delineate the four KASH-related subgroups in maize: *G* for <u>G</u>rass-specific, *S* for *At<u>S</u>INE*-like, *P* for *AtWI<u>P</u>*-like, or *T* for *AtWI<u>T</u>*-like. The genes encoding the INM LINC proteins fall into two groups, the previously published core LINC SUN domain proteins (Murphy et al., 2010), and the genes designated *MNEAP* for <u>M</u>aize *At<u>NEAP</u>*-like. Finally, the genes encoding the nucleoplasmic LINC-associated proteins are designated *NCH* for <u>N</u>MCP/<u>C</u>RWN <u>h</u>omologs or *MKAKU4* for *<u>M</u>aize <u>KAKU4</u>-like*. In summary, the 22 known or candidate core LINC and associated proteins in maize are 10 MLKs, 5 SUNs, 3 NEAPs, 2 NCHs, and 2 MKAKU4s. For all of these, the gene IDs in maizeGDB, PLAZA gene, and UniPROT are summarized in Table 1, Table S1, and diagrammed with their protein domain structures in Figure 2. Of these, the KASH and SUN proteins are the most similar to their animal counterparts, whereas the MLKGs, MLKTs, MNEAPs, MKAKU4s appear to be more characteristic of plants.

**Figure 1.**
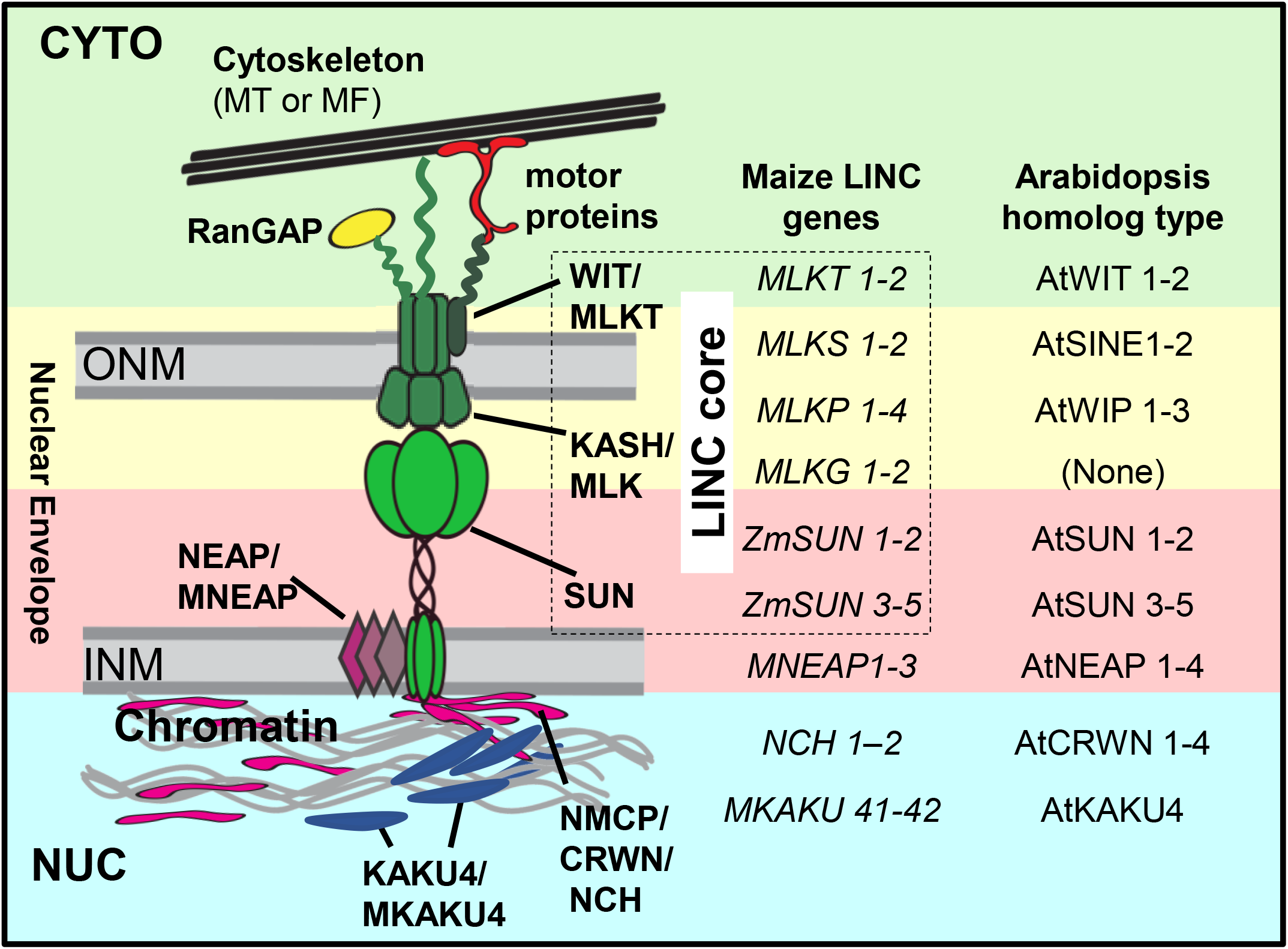
Working model of the plant LINC complex; core and associated proteins. Core LINC and associated proteins are depicted as residing in one of the four cellular locations, each represented by different background shaded colors: green, cytoplasm (CYTO); yellow, outer nuclear membrane (ONM); pink inner nuclear membrane (INM); and blue NUC (nucleoplasm). The core LINC proteins (dashed box enclosed) shown include SUN-domain proteins labeled “SUN”, KASH-related proteins “KASH/MLK”, and AtWIT-related proteins labeled “WIT/MLKT”. The LINC-associated proteins shown include cytoplasmic proteins labeled “RanGAP” and “motor proteins,” and “cytoskeleton”; the INM proteins labeled “NEAP/MNEAP”; and the nucleoplasmic proteins labeled “NMCP/CRWN/NCH” and “KAKU4/MKAKU4”. Maize gene names and their known Arabidopsis homolog types are listed. This composite model places proteins together on the basis of previously documented pair-wise interactions representing a generalized consensus assembly but not meant to depict a single complex.

**Figure 2.**
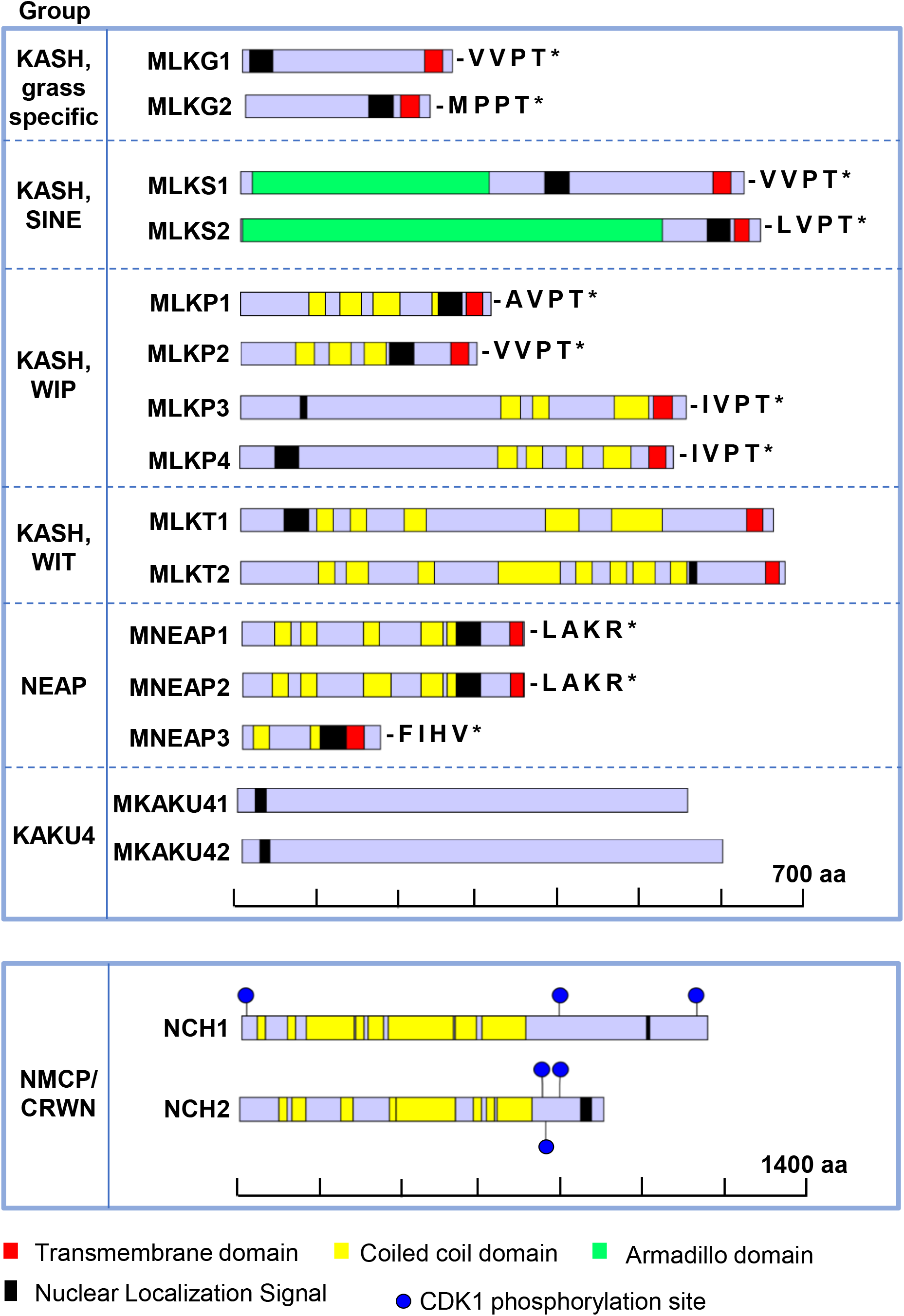
Domain organization of candidates for the maize core LINC and associated proteins complex. Schematic representations of predicted proteins from canonical transcript models are diagrammed to show the location of domains (color coded and listed at bottom). The proteins are drawn to scale, using a separate scale for the last two larger proteins, NCH1 and NCH2. The four C-terminal residues and stop codon (*) location of the KASH and MNEAP group proteins are indicated. The previously published SUN-domain proteins (Murphy et al., 2010) are not included. The partial SMC N-terminal domains in the MNEAPs are not diagrammed but are located at AA 24-308 for MNEAP1, AA 18-309 for MNEAP2, and at AA 3-109 for MNEAP3. Potential CDK1 phosphorylation sites are shown (blue circles) for NCH1 and NCH2.

**Table 1.** Maize LINC genes.

| Short Name<sup>A</sup> | Full Name<sup>A</sup> | Gene id<sup>B</sup> | Length (AA) | Domains |  |  |
| --- | --- | --- | --- | --- | --- | --- |
|  |  |  |  | Coiled coil | TMD | Other (location) |
| *MLKG1* | *Maize LINC KASH Grass-specific1* | Zm00001d002723 | 258 | 74-94 | 224-246 | KASH (247-258) |
| *MLKG2* | *Maize LINC KASH Grass-specific2* | Zm00001d038539 | 227 | - | 19-214 | KASH (215-227) |
| *MLKS1* | *Maize LINC KASH AtSINE-like1* | Zm00001d031134 | 617 | - | 579-601 | KASH (602-617) |
| *MLKS2* | *Maize LINC KASH AtSINE-like2* | Zm00001d052955 | 637 | - | 605-623 | KASH (624-637), ARM (3-517) |
| *MLKP1* | *Maize LINC KASH AtWIP-like1* | Zm00001d003334 | 308 | 82-105, 123-150, 163-197, 236-270 | 278-300 | KASH (301-308) |
| *MLKP2* | *Maize LINC KASH AtWIP-like2* | Zm00001d025667 | 290 | 68-91, 109-136, 152-179 | 258-280 | KASH (281-290) |
| *MLKP3* | *Maize LINC KASH AtWIP-like3* | Zm00001d005997 | 538 | 319-343, 358-378, 452-500 | 506-529 | KASH (530-538) |
| *MLKP4* | *Maize LINC KASH AtWIP-like4* | Zm00001d020879 | 532 | 313-337, 352-372, 401-421, 446-480 | 502-523 | KASH (524-532) |
| *MLKT1* | *Maize LINC KASH AtWIT-like1* | Zm00001d039272 | 653 | 94-114, 135-155, 201-228, 374-415, 455-517 | 620-640 | KASH (641-653) |
| *MLKT2* | *Maize LINC KASH AtWIT-like2* | Zm00001d010047 | 667 | 96-116, 130-157, 218-238, 316-392, 411-431, 453-473, 481-508, 527-547 | 643-661 | KASH (662-667) |
| *ZmSUN1* | *SUN domain protein1* | Zm00001d015450 | 462 | 169-189, 195-229 | 116-141 | SUN (277-455) |
| *ZmSUN2* | *SUN domain protein2* | Zm00001d040331 | 439 | 168-188 | 84-109 | SUN (264-478) |
| *ZmSUN3* | *SUN domain protein3* | Zm00001d042643 | 613 | 484-511 | 33-55, 554-574, 595-612 | SUN (194-358) |
| *ZmSUN4* | *SUN domain protein4* | Zm00001d012240 | 639 | 510-537 | 57-79, 580-600, 621-638 | SUN (218-382) |
| *ZmSUN5* | *SUN domain protein5* | Zm00001d011277 | 607 | 414-434, 495-523 | 46-66, 525-544, 572-588 | SUN (152-322) |
| *MNEAP1* | *Maize Nuclear Envelope-Associated Protein1* | Zm00001d010918 | 348 | 221-248 | 330-346 | NA |
| *MNEAP2* | *Maize Nuclear Envelope-Associated Protein2* | Zm00001d024756 | 347 | 149-83, 220-247 | 329-345 | NA |
| *MNEAP3* | *Maize Nuclear Envelope-Associated Protein3* | Zm00001d002875 | 170 | 14-34, 84-104 | 125-150 | NA |
| *NCH1* | *NMCP/CRWN-Homologous1* | Zm00001d051600 | 1156 | 160-278, 312-350, 362-522, 537-579, 592-701 | NA | NA |
| *NCH2* | *NMCP/CRWN-Homologous2* | Zm00001d043335 | 896 | 131-165, 250-281, 309-336, 386-532, 635-721 | NA | NA |
| *MKAKU41* | *Maize KAKU4-Like1* | Zm00001d026487 | 554 | NA | NA | NA |
| *MKAKU42* | *Maize KAKU4-Like2* | Zm00001d002012 | 591 | NA | NA | NA |

<sup>A</sup>Gene names approved by and listed at MaizeGDB (maizegdb.org)1208 <sup>B</sup>B73v4 gene models from MaizeGDB.org as of December 2017. Detailed version of this table with B73v3, PLAZA and UniPROT gene ids is provided as Table S1.

### The MLK genes: ten candidates for maize ONM core LINC genes

A hallmark of the KASH domain proteins is that they have limited sequence conservation and are not reliably recognized by sequence similarity searches. Instead, they contain a conserved structure consisting of a C-terminal transmembrane domain and a short periplasmic tail with characteristic terminal residues including a penultimate proline followed by a threonine or serine. Only a few plant genes with similarity to animal KASH genes have been identified so far (Zhou et al., 2012; Zhou et al., 2014). To find all of the potential maize LINC ONM components, we first searched the maize proteome (B73 AGPv4) from ensembl.gramene.org (Jiao et al., 2017) for KASH proteins with the following characteristic features: (1) a TM domain at the C-terminus followed by 9-40 amino acids that could extend into the perinuclear space, and (2) terminal four residues of [DTVAMPLIFY], [VAPIL], P, and T (Poulet et al., 2017a; Zhou et al., 2014). This search led to the identification of a total of eight genes encoding maize KASH-like candidates, four with homology to AtWIPs (*MLKP1* – *MLKP4*), two with homology to AtSINEs (*MLKS1* and *MLKS2*) and two found only in members of the grass family, Poaceae (*MLKG1* and *MLKG2*). We next searched the maize proteome for WIT homologs using AtWIT1 and AtWIT2 protein sequences as queries (Zhao et al., 2008) and found two maize genes, *MLKT1* and *MLKT2*, both encoding proteins with the AtWIT-like TM domains at their C-termini and multiple coiled-coil (CC) domains. Given the rapid divergence of LINC gene sequences and the lack of distinct and diagnostic domains, and gene content differences between maize cultivars (Hirsch et al., 2016), it is possible that maize has even more MLKT genes not readily detected using these criteria.

The four MLKP (AtWIP-like) genes belong to the PLAZA gene family designated HOMO4M002181, and encode proteins with multiple coiled-coil domains (MLKP1-MLKP4, Fig. 2). The previously identified AtWIP proteins interact with AtWITs and RanGAP1 and exhibit coiled-coil domain-dependent homo-and hetero-multimerization (Zhao et al., 2008; Zhou et al., 2012). The MLKP proteins are therefore predicted to interact with themselves, each other, ZmSUN proteins, and RanGAP proteins with WPP domains. The two MLKS (AtSINE-like) genes belong to PLAZA gene family designated HOMO4M002474, and encode proteins with armadillo (ARM) repeats (MLKS1-MLKS2, Fig. 2). ARM domains have been reported in a variety of proteins across plant and animal kingdoms as actin-binding and protein-protein interaction domain (Coates, 2003). The MLKS proteins are predicted, therefore, to interact and co-localize with actin, as has been demonstrated for AtSINE1 (Zhou et al., 2014). The two MLKG (grass-specific) genes lack both CC and armadillo domains and have been assigned to different PLAZA gene families - HOMO4M009187 for *MLKG1* and HOMO4M008346 for *MLKG2*.

Beyond the eight candidate MLK genes that we predict to encode maize ONM core LINC complex proteins, we also found four other maize genes that met some of our criteria. For instance, some encoded proteins with the terminal PT residues, but did not satisfy the single C-terminal TM domain criterion. It remains unknown whether these genes (B73v4 gene IDs Zm00001d024832, Zm00001d036395, Zm00001d005590 and Zm00001d017144) encode actual variant LINC proteins or simply resemble LINC proteins.

### The eight maize INM LINC genes; five SUN and three NEAP genes

The five genes encoding maize SUN-domain proteins belong to two subfamilies distinguished by the location of the SUN domain (Murphy et al., 2010). They are not newly identified in this study, but are included in Table 1 for the sake of completion, along with their most recent B73v4 gene IDs. ZmSUN1 and ZmSUN2 proteins have the canonical C-terminal SUN domain, whereas the ZmSUN3, ZmSUN4,and ZmSUN5 have a central SUN domain that is characteristic of, but not limited to plants (Graumann et al., 2014). Despite the conservation of SUN-domain proteins as a group, and their assignment to one of two subfamilies, direct assignment of orthology for individual SUN genes across taxonomic groups such as eudicots (Arabidopsis) versus monocots (maize) has yet to be established.

Another major family of INM LINC proteins are the higher plant-specific proteins called Nuclear Envelope-Associated Proteins, the NEAPs (reviewed by Groves et al., 2018; Meier et al., 2017; Poulet et al., 2017a). NEAPs exist as a multigene family, first discovered in Arabidopsis and shown to interact with SUN-domain proteins (Pawar et al., 2016). Genetic analysis of AtNEAPs showed that they affect several phenotypes, including primary root growth, nuclear morphology, and chromatin organization (Pawar et al., 2016). NEAPs have three protein structural features, the presence of extensive coiled coil domains, nuclear localization sequences, and C-terminal TM domains. We used these criteria to identify three maize NEAP genes (Table 1, Fig. 2), *MNEAP1*, *MNEAP2*, and *MNEAP3*. Interestingly, we observed partial homology to N-terminus of structural maintenance of chromosomes domain (SMC_N, PFAMID PF02463) proteins in the MNEAPs, further implicating them in DNA and chromosomal functions.

The MNEAP1 and MNEAP2 proteins both end with a characteristic C-terminal hydrophobic patch and two basic terminal residues KR (Pawar et al., 2016). These two proteins share 92% sequence identity and are very similar to the protein encoded by one of the two sorghum NEAPs genes (sobic.009G033800). The *MNEAP1* and *MNEAP2* most likely arose from the genome duplication event that occurred following the divergence of maize from sorghum, ~ 11 MYA (Gaut et al., 2000). *MNEAP3* fits our initial criteria but is otherwise quite different in many respects including an unusually large number of splice variant transcript models and the lack of KR terminal residues. Consequently, *MNEAP3* may be a complex locus or alternatively, pseudogene no longer under evolutionary pressure to maintain its structure.

### The four maize nucleoplasmic LINC-associated genes; two *NCH* and two *KAKU4* genes

In mammals, the SUN-domain proteins of the LINC complex interact with the nuclear lamina, which is located at the periphery of the nucleoplasm and made up of lamin proteins. Plant nuclei lack laminA/B orthologs, but they do have a lamin-like meshwork that is thought to consist of lamin-like proteins known as NMCP/CRWN as well as CRWN-binding nuclear proteins, KAKU4, (Ciska et al., 2013; Goto et al., 2014). These proteins collectively are thought to comprise a significant portion of the plant nucleoskeleton and they are coupled to the LINC complex via interactions with SUN domain proteins (Goto et al., 2014).

Using AtCRWN and DcNMCP protein sequences as queries, we identified two maize genes, *NCH1* and *NCH2*, representing the two major clades of NMCP/CRWN genes (Ciska et al., 2013; Wang et al., 2013). The maize *NHC1* is most similar to *DcNMCP1*, *DcNMCP3*, and *AtCRWN1*. The genes in this group are members of PLAZA gene family HOM04M004640 (Table S1). The maize *NCH2* gene is most similar to *DcNMCP2*, *AtCRWN2*, *AtCRWN3*, and *AtCRWN4*. The genes in this group are members of the PLAZA gene family HOM04M003564. Both NCH1 and NCH2 proteins share several features with their plant homologs. These features include large proteins (~100 kDa or greater), multiple coiled-coil domains over the majority of the protein, multiple CDK1 phosphorylation sites, and a nuclear localization sequence near the C-terminus (Fig. 2).

Using AtKAKU4 protein sequence similarity searches, we identified two highly similar maize KAKU4-like genes, *MKAKU41* and *MKAKU42*. These two genes and AtKAKU4 are all members of the PLAZA gene family HOM04M004991. KAKU4 and KAKU4-like genes appear to be limited to flowering plants and encode proteins that have been described as candidate lamina components (Goto et al., 2014). As a group, they may be remotely associated with the LINC complexes but are included here because of the known interaction of AtKAKU4 with AtCRWN proteins as well as the genetic evidence that implicates them in LINC functions and nuclear architecture (Goto et al., 2014). Aside from a NLS near the N-terminus, the MKAKU4 proteins do not contain any notable structural protein domains or motifs (Fig. 2).

### A mix of ancient and derived genes encode components of the maize LINC complex

Given that the time of appearance and the evolutionary conservation of genes often reflects functional properties, we examined the maize LINC candidate genes and their counterparts in a variety of plant species as summarized in Figure 3. Recent phylogenetic analyses of plant LINC genes or sub-families therein have been published (Ciska et al., 2013; Poulet et al., 2017a; Zhou et al., 2014), with emphasis on Arabidopsis and other eudicot species, but did not include the latest PLAZA gene families or maize B73v4 annotations (Jiao et al., 2017; Van Bel et al., 2018). The species listed (Fig. 3) are arranged in increasing evolutionary distance from maize starting with sorghum and extending to the unicellular green alga, *Chlamydomonas reinhardtii*.

In plants, the most highly conserved members of the core LINC and associated complex belong to gene families that encode the C-terminal and mid-SUNs, NCH2 and the AtSINE-like KASH proteins MLKS1 and MLKS2). These gene families are present in all the plant species listed (shaded blue or green, Fig. 3). Curiously, of all the genes listed, only the mid-SUN genes are listed as present in Chlamydomonas PLAZA gene families. The next most highly conserved plant LINC gene families encode the MNEAP and NCH1 proteins. They are present in seed-bearing plants (shaded tan, Fig. 3), which includes pine, a conifer species. The third most highly conserved plant LINC gene families encode MLKP, MLKT and MKAKU4 proteins, all of which are found in flowering plants, the angiosperms (shaded pink, Fig. 3). In contrast, the most recent to appear in evolutionary time are the gene families that encode the MLKG proteins, present in the grass family within the monocots (shaded yellow, Fig. 3). The biological significance of the timing of evolutionary appearance remains to be determined, but we generally expect that the more widely distributed gene families have more ancient or basal functions, whereas the more recent gene families could have resulted from duplication followed by either regulatory divergence, such as tissue-specific expression, or gain of new biological functions.

**Figure 3.**
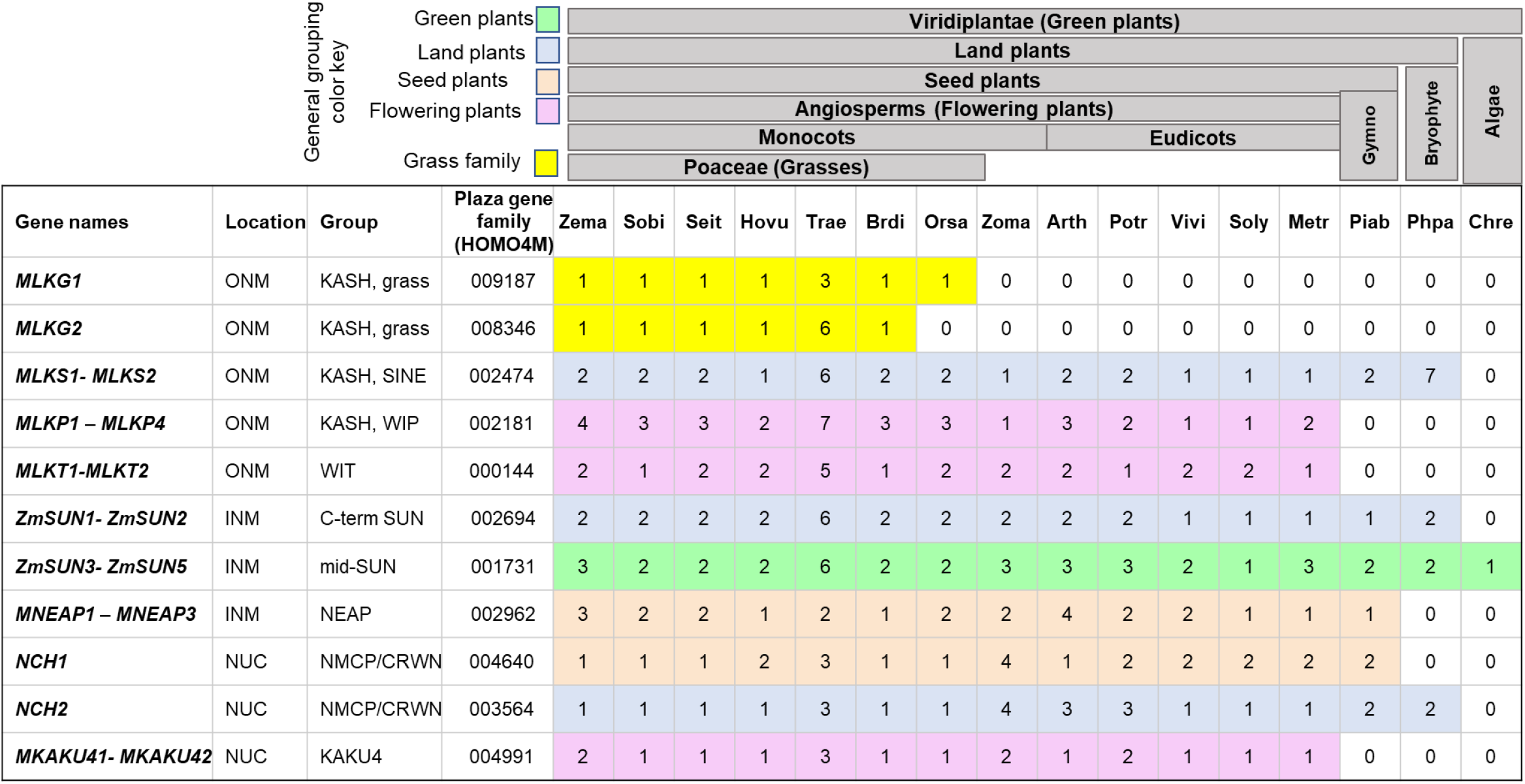
Evolutionary conservation of plant LINC complex components. Candidate maize LINC gene names, cellular location predictions, gene family group, and PLAZA gene family ID (each starting with shared prefix of HOMO4M) are listed in the first four columns. The other columns provide counts of the numbers of genes for each species (top row) in each Plaza gene family (4th column). The general grouping color key is indicated on the top left of the figure. Species included are Zema, *Zea mays;* Sobi*, Sorghum bicolor,* Seit, *Setaria viridis;* Hovu, *Hordeum vulgare;* Trae,*Triticum aestivum;* Brdi, *Brachypodium distachyon;* Orsa*, Oryza sativa spp. Japonica;* Zoma*, Zostera marina;* Arth*, Arabidopsis thaliana;* Potr*, Populus trichocarpa;* Vivi*, Vitis vinifera;* Metr*, Medicago truncatula;* Piab*, Picea abies;* Phpa*, Physcomitrella patens* and Chre, *Chlamydomonas reinhardtii.* Broader grouping names are provided along the top (labeled grey boxes).

### Maize LINC genes exhibit tissue-specific and developmental co-expression patterns

We analyzed the expression patterns of the maize LINC candidate genes as shown in Figure 4 in order to examine them for evidence of expression and potential functional groupings that should be reflected in their differential or co-expression patterns. For this, we used the expanded maize gene expression atlas which contains transcriptome data from numerous tissues broadly sampled across most of the developmental stages of the maize life cycle (Stelpflug et al., 2016). For comparison, we included three housekeeping genes (HKG, Fig. 4A), *CDK1*, *RPN*, and *SGT1* (Lin et al., 2014). We excluded *MNEAP3*, which lacked a gene model when the atlas was produced, and *MLKG1*, which had zero reads in all fifteen tissues selected for this study. Despite their exclusion, these two are presumed to be real genes and not pseudogenes because of the expression of their corresponding transcripts detected in one or more tissues (Chettoor et al., 2014; Walley et al., 2016; Wang et al., 2016; Xu et al., 2012).

**Figure 4:**
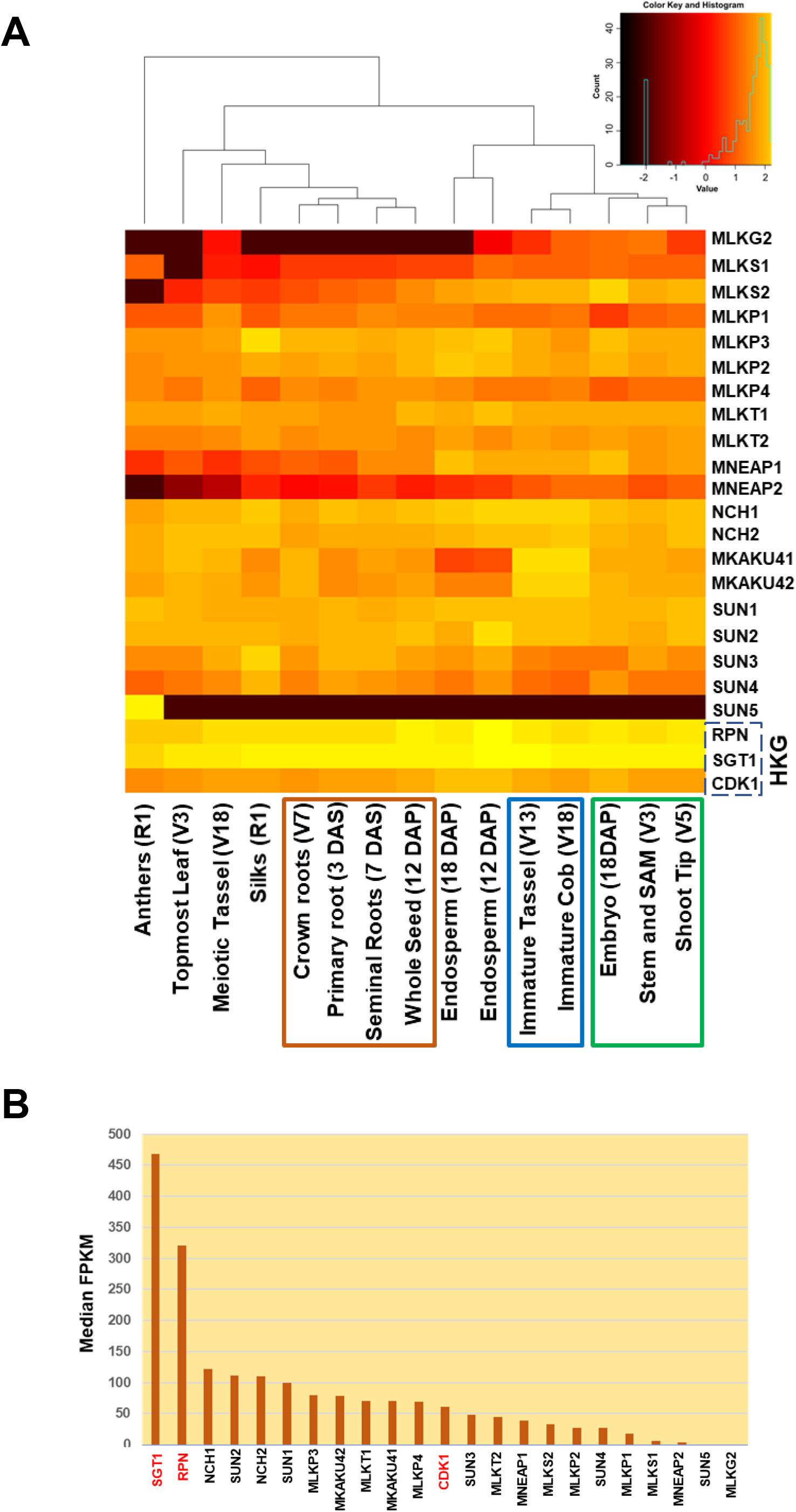
Gene expression levels and co-expression clustering of 20 maize LINC genes. Expression levels of 20 maize candidate LINC genes were analyzed using published transcriptome data (Stelpflug et al., 2016) and including the genes designated as housekeeping genes (HKG) in a previous study (Lin et al., 2014). (A) Expression profiles of LINC genes in several tissues developmental stages. Data are presented as log10 Fragments Per Kilobase of exon per Million fragments mapped (FPKM) values from published transcriptome. Hierarchical clustering grouped tissues according to similar expression profiles. The root, young inflorescence, and meristem groups are denoted by orange, blue, and green boxes, respectively. The growth stage of maize plant at the time of tissue harvesting are shown in parentheses; V, vegetative stage; R, reproductive stage; DAP, days after pollination. (B) The Median FPKM values of the tissues shown in panel A are plotted, ranked by value for individual genes. Housekeeping genes are shown in red.

Most of the candidate maize core LINC and LINC-associated genes showed a pattern of broad expression across multiple tissues, with co-expression patterns (dendrogram, Fig. 4A) revealing several distinct groupings. The “root” co-expression group (orange box, Fig. 4A) includes three root samples and 12 DAP developing seed. The “young inflorescence” co-expression group (blue box, Fig. 4A) includes male (immature tassel) and female (immature cob) tissues. The “meristem” co-expression group (green box, Fig. 4A) includes the developing embryo and growing shoot tip tissues. These last two groups together form a larger co-expression cluster from tissues that share the property of having actively dividing cell populations. Beyond these three groups, several tissues show distinctive patterns of LINC gene expression, not clustering with other tissues, including anthers (male pollen-bearing organs), topmost seedling leaf, meiotic tassel, and silks (female, pollen-receiving organs).

To characterize the general expression levels of the candidate maize LINC genes, we plotted the median FPKM values across different tissues (Fig. 4B). Maize LINC genes generally exhibit low to moderate expression levels, similar to those of the house-keeping *CDK1* gene (Fig. 4B). The more highly expressed LINC-related genes had transcript levels that ranged from 80-140 median FPKM; these genes included the MKAKU4s, the C-terminal SUNs, the NCHs, and some of the KASH genes, *MLKP3* and *MLKP4*. The less-expressed genes had transcript levels that ranged from 3-50 median FPKM; these included the mid-SUNs, the MNEAPs, and most of the other KASH genes. As a group, the maize AtWIP-like genes (*MLKP1-4*) have higher expression levels than the AtSINE-like genes (*MLKS1*, *MLKS2*), with *MLKP3* and *MLKP4* showing the most uniform and broad expression. In contrast, the mid-SUN *SUN5* gene exhibited a most striking pattern of expression in only pollen. The *Poaceae*-specific gene, *MLKG2,* appears to be expressed in tissues characterized by meristem and active cell division, but absent from others such as endosperm, root, leaf, and silks.

Among the highly similar gene pairs that are likely to have arisen as part of a recent genome duplication (Gaut et al., 2000), some of them (*MLKP1 and MKLP2; MLKP3 and MLKP4; and MKAKU41 and MKAKU42*) show similar expression levels whereas other gene pairs, such as *MNEAP1* and *MNEAP2,* exhibit differential expression levels. Overall, this expression analysis provides evidence that genes encoding LINC and associated proteins are transcribed and regulated in a tissue-specific manner. The co-expression groupings predict possible but untested genetic or physical interactions that may reflect shared functions.

### Identification of LINC and associated proteins from endogenous maize co-IP using anti-ZmSUN2 antibodies

In order to find naturally occuring interactors of maize SUN proteins, we carried out co-immunoprecipitation (co-IP) optimized for NE proteins from purified earshoot nuclei as summarized Table 2. Maize earshoots of ~3-5 cm in length yield high quality cellular extracts from fast-growing somatic tissues several days prior to silking and fertilization. The antibodies used for co-IP were the affinity-purified serum designated anti-SUN2-NPAP as previously characterized (Murphy et al., 2014). Control co-IP experiments were conducted using either no antibodies or irrelevant but similarly affinity-purified rabbit sera, anti-ZmNDPK1 (Kopylov et al., 2015). A key technical advance was the use of the short-chain crosslinker dithiobis(succinimidyl propionate), DSP. We adapted the membrane protein cross-link immunoprecipitation (MCLIP) method for use with maize because it was shown to effectively recover SUN-interacting proteins from U2OS cells (Jafferali et al., 2014). We found that the MCLIP method, using DSP together with light formaldehyde fixation to preserve the nuclei, resulted in substantially increased final recovery and detection of known or predicted maize homologs of plant LINC, NE, or SUN-interacting proteins (Tamura et al., 2013; Wang et al., 2013; Zhou et al., 2012).

**Table 2.** Proteins co-precipitated with ZmSUN2 pull down on maize ear shoot nuclei.

|  | Maize gene id (B73v4) | Maize gene name | Gene Description | Exclusive unique peptide count (IP46) | SUN IP (Normalized total spectra) | Irrelevant antibody control IP | No Antibody control IP |
| --- | --- | --- | --- | --- | --- | --- | --- |
| **Core LINC components** | Zm00001d040331 | ZmSUN2 | SUN domain protein2 | 11 | 384.45 | 0 | 0 |
|  | Zm00001d015450 | ZmSUN1 | SUN domain protein1 | 3 | 78.398 | 0 | 0 |
|  | Zm00001d051600 | NCH1 | Protein CROWDED NUCLEI 1 | 2 | 82.803 | 0 | 0 |
|  | Zm00001d043335 | NCH2 | Nuclear lamina component | 3 | 28.871 | 0 | 0 |
|  | Zm00001d002723 | MLKG1 | OSJNBa0010H02.21-like protein | 2 | 14.459 | 0 | 0 |
|  | Zm00001d052955 | MLKS2 | ARM repeat superfamily protein | 2 | 1.2886 | 0 | 0 |
|  | Zm00001d010047 | MLKT2 | WPP domain-interacting tail-anchored protein 1 | 1 | 26.117 | 0 | 0 |
| **Nucleoporins** | Zm00001d052282 | NUA | Nuclear-pore anchor | 7 | 87.161 | 0 | 0 |
|  | Zm00001d048023 | NUP133 | Nuclear pore complex protein NUP133 | 1 | 18.911 | 0 | 0 |
|  | Zm00001d046566 | NUP96 | Nuclear pore complex protein NUP96 | 1 | 7.253 | 0 | 0 |
|  | Zm00001d026263 | NUP88 | Nuclear pore complex protein NUP88 | 1 | 7.206 | 0 | 0 |
|  | Zm00001d005355 | GP210 | Nuclear pore complex protein GP210 | 1 | 7.206 | 0 | 0 |
| **Cytoskeletal elements** | Zm00001d013410 | Actin-1 | ARP7 Nuclear actin | 3 | 82.756 | 0 | 0 |
|  | Zm00001d012556 | Tub6 | beta tubulin6 | 5 | 40.576 | 0 | 0 |
|  | Zm00001d015348 | Tub4 | Tubulin beta-4 chain | 1 | 7.253 | 0 | 0 |
| **Ribosomal proteins** | Zm00001d044287 | L19 | Ribosomal protein L19 | 2 | 40.623 | 0 | 0 |
|  | Zm00001d017047 | L6 | 60S ribosomal protein L6 | 1 | 30.569 | 0 | 0 |
|  | Zm00001d022463 | L22-2 | 60S ribosomal protein L22-2 | 1 | 11.658 | 0 | 0 |
| **RAN** | Zm00001d017239 | RanGAP2 | RAN GTPase-activating protein 2 | 1 | 14.459 | 0 | 0 |
|  | Zm00001d048223 | RABA6a | Ras-related protein RABA6a | 2 | 14.412 | 0 | 0 |
| **DNA/RNA binding** | Zm00001d052820 | RNA binding protein | RNA-binding (RRM/RBD/RNP motifs) family protein | 2 | 21.665 | 0 | 0 |
|  | Zm00001d025638 | bZIP5 | Basic-leucine zipper domain | 1 | 11.658 | 0 | 0 |
|  | Zm00001d042267 | ARF10 | Auxin response factor 10 | 1 | 11.658 | 0 | 0 |
|  | Zm00001d011297 | MYBR35 | Putative MYB DNA-binding domain superfamily protein | 1 | 7.206 | 0 | 0 |
|  | Zm00001d016354 | bHLH72/bHLH041 | Putative transcription factor bHLH041 | 1 | 7.206 | 0 | 0 |
| **Histone** | Zm00001d034479 | H1a | Histone 1 (H1) | 1 | 26.117 | 0 | 0 |

The proteins recovered from the co-IP assays were subjected to tryptic digestion and identified by liquid chromatography tandem mass spectrometry (LC-MS/MS). Over 140 proteins were identified (Table S3) using software thresholds of 95% protein and 95% peptide. As expected, SUN2 was the most abundant protein, but we also repeatedly detected SUN1, which could result from cross-reactivity of ZmSUN1 with our anti-SUN2 serum, or from actual co-IP of ZmSUN1 bound to SUN2, as expected from documented hetero multimerization of SUN proteins in plants and animals (Sosa et al., 2012).

The identified co-precipitated proteins were considered high confidence if they were detected in multiple replicates but not in the negative controls. The specificity of the modified MCLIP method is evidenced by recovery of core LINC components and predicted cytoplasmic and nucleoplasmic proteins. Among the predicted core LINC components repeatedly found in the top co-IP hits were three ONM proteins (MLKG1, MLKS2, and MLKT2) two INM proteins (ZmSUN1 and ZmSUN2), two NMCP/CRWN homologs (NCH1 and NCH2), two RAN family proteins (RAN GTPase and RAN-GAP2) and five nuclear pore proteins (NUP133, NUP96, NUP88, GP210, and NUA).

We also found proteins that were co-precipitated but likely via indirect interaction with SUN. These include the cytoskeletal proteins TUB6, TUB4 and Actin-1/ARP7. The Actin-1/ARP7 protein could be cytoplasmic or nucleoplasmic or both because its annotation includes the designation of nuclear actin, an intriguing possibility in light of recent findings showing cross-species binding of Arabidopsis AtARP7 to the carrot nuclear matrix protein DcNMCP1 (Mochizuki et al., 2017). We recovered several nuclear proteins with DNA-binding activities including transcription factors bZIP, ARF10, and the linker histone H1a. Overall, the co-IP resulted in confirmatory identification of naturally occuring SUN2-interacting proteins with known or suspected LINC proteins, plus several additional proteins that represent new targets for future investigation. To further verify some of the LINC candidates from either the Co-IP or bioinformatic analyses, we selected three genes for further localization and interaction assays *in vivo* using transient heterologous expression assays.

### *In planta* evidence of NE targeting and ZmSUN2 interaction for three candidate maize LINC proteins

In order to investigate the subcellular localization of the maize KASH proteins, we transiently expressed N-terminal GFP fusions of MLKG1, MLKP1 or MLKP2 in *Nicotiana benthamiana* leaf tissue as summarized in Figure 5. Expression of a positive control, mCherry-ZmSUN2, shows that maize NE proteins can be properly expressed from the plasmids and localized to the NE in this leaf cell heterologous expression system. All three of the previously untested ONM maize LINC candidate proteins, MLKG1, MLKP1 and MLKP2, also localized to the nuclear periphery (Fig. 5A). These localization patterns are comparable to those previously reported for AtKASH proteins in similarly designed expression assays (Zhou et al., 2012; Zhou et al., 2014). For GFP-MLKG1, a network-like pattern at the cell periphery was also seen (Fig. 5B; Fig. S2C). In order to further examine this pattern, we co-expressed GFP-MLKG1 with an ER marker and an actin marker and found that GFP-MLKG1 co-localizes with both the ER and the actin network (Fig. 5B). Similar co-localization patterns have previously been demonstrated for AtSINE1 (Zhou et al., 2014) but, unlike AtSINE1, MLKG1 does not possess an ARM domain or other known actin binding motifs.

**Figure 5.**
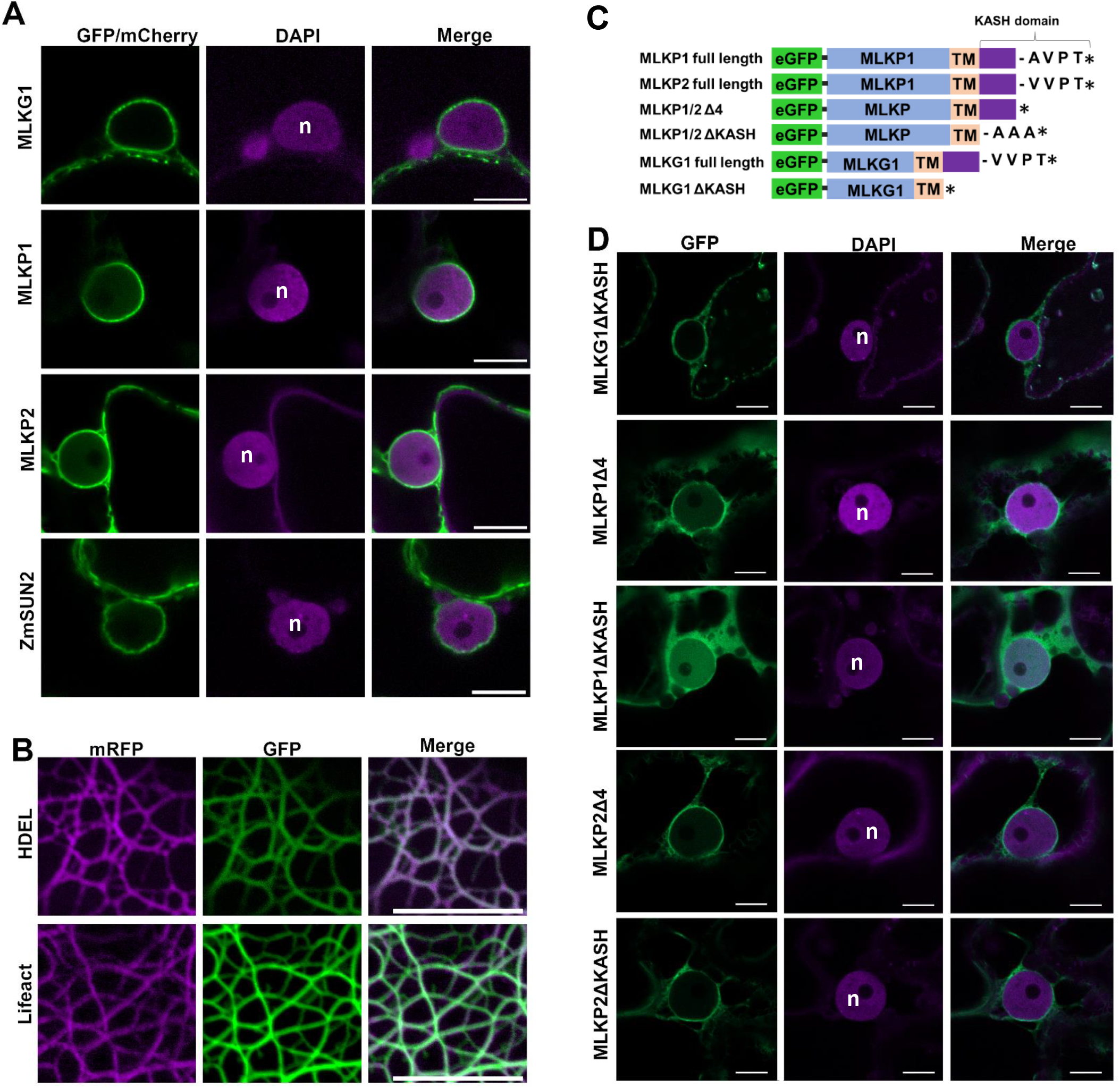
Subcellular localization of transiently expressed GFP-MLKP fusions in *N. benthamiana*. Agrobacterium-mediated leaf transformation with expression constructs was carried out followed by imaging three days later using 100X lens with confocal microscopy to detect DAPI (405 nm excitation laser), GFP (488 nm excitation laser), or mCherry/mRFP (561 nm excitation laser). Representative focal sections are shown for GFP (green), mCherry (green in panel A, bottom row) mRFP (majenta in panel B), or DAPI (majenta in panels A and D). (A) GFP-MLKG1, GFP-MLKP1, GFP-MLKP2 and mCherry-ZmSUN2 (green) are localized at the nuclear periphery, surrounding the nucleus (n) stained with DAPI (magenta). (B) GFP-MLKG1 (green) is also present in a network-like pattern at the cell periphery, where it partially co-localizes with the ER marker HDEL-mRFP (magenta, top row) and the actin marker lifeact-mRFP (magenta, bottom row). (C) Schematic overview depicting the terminal deletion constructs. (D) Images from deletion constructs (listed at left) are shown as GFP only (1st column), DAPI only (2nd column), or overlay (3rd column). GFP-MLKP1ΔKASH shows additional localization to the cytoplasm and the nucleoplasm (magenta), whereas the others retain a tendency to localize to the nuclear periphery. All scale bars 10μm.

The C-termini of the KASH proteins are known to play a key role in its NE localization which is mediated through interaction with SUN proteins (Sosa et al., 2012; Zhou et al., 2012; Zhou et al., 2014). To test for a role of the C-terminal residues in our maize KASH candidates, we created and tested several classes of C-terminal deletion constructs (Fig. 5C). The Δ4 deletion constructs are lacking the last 4 C-terminal amino acids. The ΔKASH deletion constructs are lacking all of the amino acid residues following the TM domain, made with (MLKP1/2 ΔKASH) or without (MLKG1ΔKASH) a minor three alanine stabilizing spacer. The ΔTM deletion constructs are lacking the entire C-terminus starting from the beginning of the predicted TM domain (Fig. 5C, S1,S2). For MLKP1, both the ΔKASH or ΔTM protein truncations changed the localization pattern to that resembling soluble proteins (Fig. 5D, S2) whereas the Δ4 truncation showed a small increase in apparent solubility, but retained considerable targeting to the NE. For MLKP2Δ4, MLKP2ΔKASH, and MKLG1ΔKASH, the terminal truncations still resembled that of the full length proteins with localization to the NE. For comparison, we expressed GFP-only as a control (Fig. S2B), which illustrates the distribution of a soluble protein without a TM domain in the cytoplasm and nucleoplasm. Interestingly, the removal of the KASH domains from either MLKP2 or MLKG1 did not abolish its ability to localize to the nuclear periphery, suggestive of other or additional determinants that enable MLKP2 and MLKG1 associations with the NE membranes (e.g. Fig. 5B, S2C).

In order to investigate whether maize KASH binds to maize SUN *in vivo*, we co-expressed three different maize GFP-KASH with or without ZmSUN2. The GFP-KASH proteins co-localized to the NE with mCherry-ZmSUN2 full length (Fig. S7). If GFP-KASH and ZmSUN2 do interact, we would expect to see a reduction in the mobility of GFP-KASH as measured by fluorescence recovery after photobleaching and indicative of interacting partners, as previously described (Zhou et al., 2014). For MLKG1 and MLKP2, we did indeed observe reduced mobility when co-expressed with ZmSUN2 (Fig. 6B,F; S4-5). When GFP-MLKG1 is transiently expressed by itself, 63±8% of the protein population is mobile while 36% remains immobilized. However, upon co-expression with mCherry-ZmSUN2, only 49±12% of the protein population was mobile, indicating that the co-expressed ZmSUN2 substantially increased the immobile fraction of GFP-MLKG1 from 36% to 51% (Fig. 6A,B). We interpret this shift as indirect evidence for ZmSUN2 interaction with GFP-MLKG1. To test this idea, GFP-MLKG1 was co-expressed with a domain deletion mutant of ZmSUN2 lacking the SUN domain (mCherry-ZmSUN2ΔSUN); we found that the mobility of MLKG1 was similar to that of the MLKG1 alone (68±7%; Fig. 6 B,C). This finding indicates that the SUN domain is required for mediating the change in GFP-MLKG1 mobility, providing strong indirect evidence that ZmSUN2 interacts with MLKG1. To further test if the coiled-coil domain of the SUN protein is required for reduced mobility, we created a version of mCherry-ZmSUN2 lacking the coiled-coil domain, mCherry-ZmSUN2ΔCC. Co-expression of this mCherry-ZmSUN2ΔCC with GFP-MLKG1 had a similar effect on KASH mobility to that of the full length SUN2 (Fig. 6 B, C; S4-5). This result establishes that the ZmSUN2-co-expression-dependent increase in the immobile fraction of MLKG1 occurs to a similar degree when co-expressed with ZmSUN2 with or without the CC domain.

**Figure 6.**
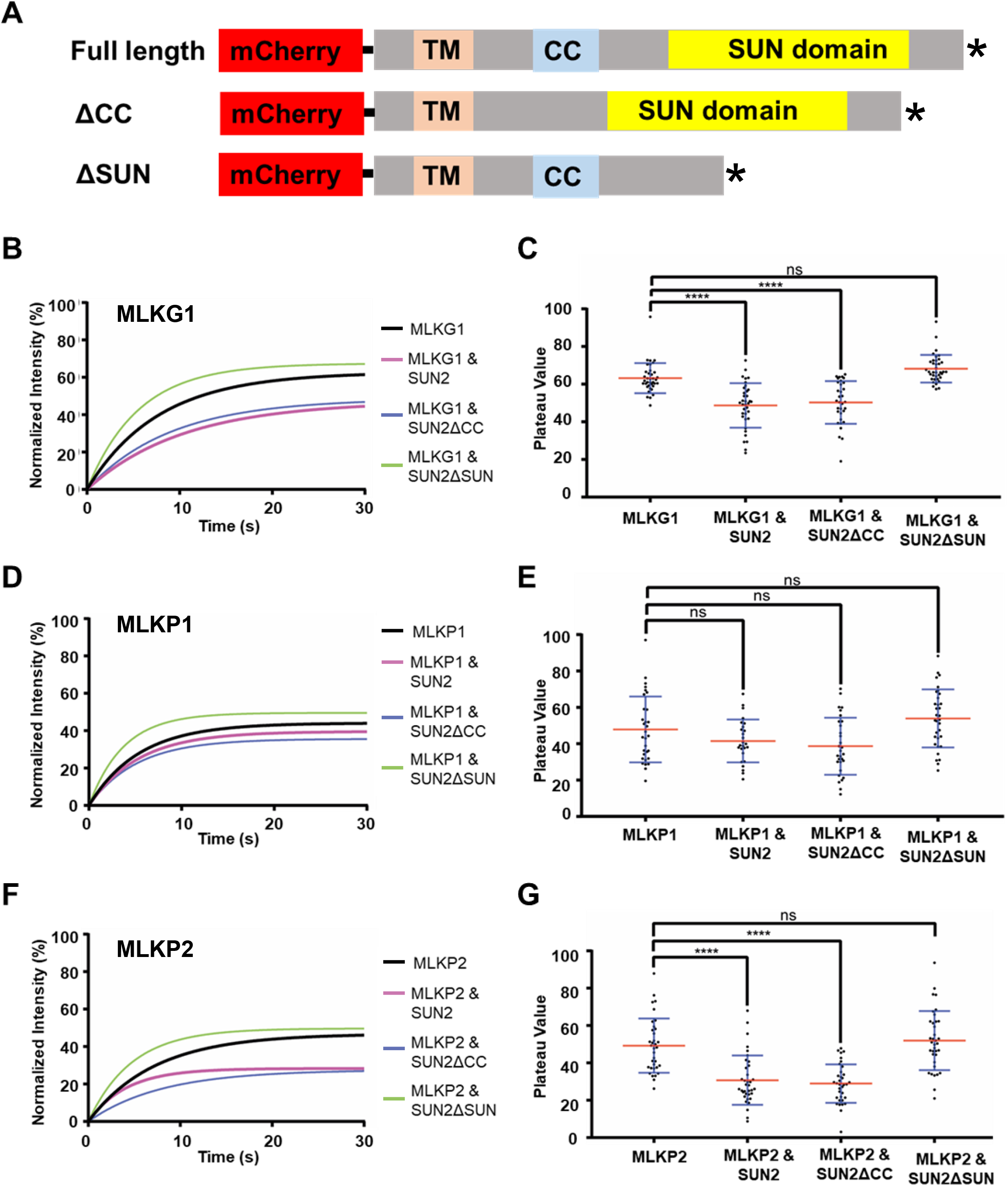
Fluorescence recovery after photobleaching (FRAP) co-expression interaction assays of MLK LINC candidates with ZmSUN2 in *N. benthamiana*. (A) Schematic overview depicting the full length mCherry-ZmSUN2 and internal deletion constructs that removed the coiled coil domain (ΔCC) or the SUN domain (ΔSUN). Normalized averaged (n=26-37) intensity FRAP curves for (B) GFP-MLKG1, (D) GFP-MKLP1, or (F) GFP-MLKP2 expressed alone (green), with full-length mCherry-SUN2 (black), with mCherry-SUN2-ΔCC (blue), or with mCherry-SUN2-ΔSUN (pink). Plateau values are plotted to the right for assays with GFP-MLKG1 (C), GFP-MLKP1 (E), and GFP-MLKP2 (G). Box plot error bars represent SD (blue) and mean (red) values. ANOVA statistical analyses with multiple comparisons are noted (ns = not significant = p≥0.05, **** = significant with P ≤0.0001).

Similar changes in protein mobility were observed for GFP-MLKP2 (Figs 6 F, G and S4-5) when co-expressed with full-length ZmSUN2, ZmSUN2ΔSUN or ZmSUN2ΔCC. Co-expression with ZmSUN2 or ZmSUNΔCC resulted in significant increases in the immobile fraction of GFP-KASH from 51% to 69% or 71%, respectively. In contrast, we did not observe statistically significant changes in GFP-MLKP1 mobility when co-expressed with full-length or deletion constructs of ZmSUN2 (Figs 6D, E and S4), even though the changes observed were all in the same directions as those for GFP-MLKG1 and GFP-MLKP2 (Figs 6, S3-5).

Of the 3 candidates tested, MLKG1 was already found by endogenous maize co-IP to interact with SUN2, but the other two, MLKP1 and MLKP2 were not, presumably because of their low expression levels in earshoot (Fig. 4). In order to check for their ability to interact with SUN2 *in planta*, we assayed them by co-IP in tobacco tissues where they were co-expressed as heterologous fluorescent fusion proteins (Fig. S6A). Using anti-GFP antisera, we observed co-precipitation of mCherry-SUN2, but not in control extracts from non-infiltrated or mCherry-SUN2 only samples. These additional tests confirmed that all three of the proteins tested, MLKG1, MLKP1, and MLKP2, showed SUN2 interaction in one or more direct or indirect interaction assays (Table in Supplemental Figure 6B).

In summary, all three of the KASH candidates tested showed NE localization in the transient heterologous expression system. They also showed evidence of in planta interaction with ZmSUN2, supported by one or more lines of evidence, co-expression FRAP, endogenous co-IP, or heterologous co-IP. These findings, together with our bioinformatic, transcriptomic, and phylogenetic analyses provide the first experimental evidence for an entire core LINC complex and associated proteins in a model monocot species.

## DISCUSSION

The human population growth rate imposes a great challenge in agriculture and food production. It has been estimated that in order to provide food for a projected population of 9.15 billion in 2050, agricultural productivity must increase by 60% (from 2013 Summit Report at https://plantsummit.wordpress.com/). For both biological and social reasons, it is necessary to discover and characterize new plant genes that inform our knowledge how the cell nucleus impacts organismal traits. This is especially true for model crop species such as maize with annual production at over 400 million bushels per year (FAOSTAT 2016, http://www.fao.org/faostat/en/). This study, the first to define the composition of the maize LINC complex and associated proteins, represents a major advance that leverages knowledge from non-plant eukaryotes regarding the role of the LINC complex in gene regulation, chromatin dynamics, and nuclear architecture (reviewed by Alam et al., 2016; Hieda, 2017; Kim et al., 2015; Poulet et al., 2017b).

Here we set out to systematically identify all of the maize LINC core and associated components, adding a valuable new and different model species for investigation of the plant nuclear envelope and important biological processes such as meiotic chromosome behavior, development, signaling, and pathology (Kracklauer et al., 2013; Meier et al., 2017; Razafsky and Hodzic, 2015; Tapley and Starr, 2013). Maize provides a valuable plant complement to Arabidopsis. Both maize, a monocot, and Arabidopsis, a eudicot, have made important contributions to biology, yet they have quite different body plans, seed development pathways, genome sizes, and gene expression profiles, having diverged 200 MYA (Brendel et al., 2002; Conklin et al., 2018; Dukowic-Schulze et al., 2014; Wang et al., 2012; Zeng et al., 2014).

In Arabidopsis, SUN1 and SUN2 double knockouts are known to disrupt conserved functions such as meiotic progression, excessive chromosomal interlocks, and reduced fertility (Varas et al., 2015). However, Arabidopsis does not exhibit a canonical early prophase telomere bouquet cluster on the nuclear envelope, nor has it been shown to have a meiotic SUN belt (Armstrong et al., 2001; Hurel et al., 2018; Murphy et al., 2014). These differences may highlight the limitations of extrapolating across divergent species within the plant kingdom. Furthermore, the difficulty in identifying certain orthologs for some LINC components illustrates the importance of defining the composition of the LINC complex in multiple species, especially in species with well-developed genetic and cytological resources that enable comparative analysis of the associated biological functions.

Searching for maize KASH and associated ONM proteins, we found homologs of plant genes encoding AtWIP and AtSINE1/2-like proteins, but not those encoding AtSINE3/4, SINE5 or AtTIK (Graumann et al., 2014; Zhou et al., 2014). Curiously, the *AtSINE3/4* genes are found only in brassicaceae, the *SINE5* genes are found only in medicago, and the *AtTIK* genes are found only in Arabidopsis (Poulet et al., 2017a). Similarly, *MLKG1* and *MLKG2* genes are found only in maize or other grasses and appear to be completely absent from eudicot species such as Arabidopsis or Medicago. In contrast, the *MLKT* genes of maize are clearly homologous to the *AtWITs* genes, a rare case in which sequence conservation is sufficient to identify candidate ONM LINC components. Notably, additional maize genes resembling ONM LINC components were found and may have NE functions, but fell short of the criteria we applied in this study. It remains an intriguing but recurring observation that ONM LINC proteins are part of an ancient structure, but they can be highly divergent. In this regard, plant KASH genes are similar to those from opisthokonts, with characteristic variation in protein sequence and structure (Meier et al., 2017).

Searching for additional maize genes encoding INM proteins, we found three homologs of the plant-specific *NEAP gene*s. The *NEAP* genes are limited to seed plants (Poulet et al., 2017a) and not found in any of the representatives of the more ancient moss and algal species included in this study (Fig. 3). The *MNEAP1* and *MNEAP2* genes encode protein sequences that end in two basic residues, like their eudicot homologs, but the functional significance of these terminal residues remains unknown. Interestingly, the maize NEAPs have SMC_N domains which are currently uncharacterized but may point to a chromosome tethering or nuclear structural role. The maize NEAPs named in this study are three of the eleven maize genes assigned to the PLAZA gene family HOM04M002962. Consequently, the actual number of the maize NEAP genes may grow as revised bioinformatic and functional criteria for their recognition emerges. In contrast, we did not find evidence for additional SUN genes of maize beyond the five previously described (Murphy et al., 2010; Murphy et al., 2014).

Searching for nucleoplasmic proteins associated with plant LINC complexes, we found two maize genes with homology to *NMCP*/*CRWN* genes, *NCH1* and *NCH2*, both of which are deeply conserved in plants. Each of these are associated with multiple transcript models (Wang et al., 2016) suggestive of splice variants, an intriguing observation in light of the tendency for animal lamin genes to also have many splice variants (Broers et al., 2006). Identification of plant nuclear matrix components that interact with the nuclear envelope provides a more complete view of the physical bridge across the NE. As such, this emerging model provides new opportunities to investigate the mechanisms that underlie how signals are transduced via the LINC complex.

Cellular localization experiments confirmed the NE localization of fluorescent fusion protein constructs for ZmSUN2 and all three of the maize KASH homologues tested, MLKG1, MLKP1 and MLKP2 (Fig. 5). Interestingly, GFP-MLKG1 co-localizes with the ER and the actin network, implicating it in cytoskeletal signal transduction to the nucleus. Relatedly, Arabidopsis actin is tethered to the LINC complex through direct interactions with AtSINE1 and through the interaction of myosin XI-i with WIT-WIP (Tamura et al., 2013; Xu et al., 2007; Zhou et al., 2014). The presence of *SINE1*, *WIP* and *WIT* homologs in maize, together with the poaceae-specific *MLKG1,* suggests that plant genomes may encode a diverse repertoire of ONM proteins to connect and integrate NE with cytoskeletal processes.

To provide additional experimental evidence on the composition of the maize LINC complex, we used a ZmSUN2 co-IP method optimized for membrane proteins together with isolated nuclei. We found a large number of different proteins that were generally consistent with predictions from our working model (Fig. 1) and previous reports (Dittmer and Richards, 2008; Goto et al., 2014; Pawar et al., 2016; Tamura et al., 2010; Tamura et al., 2013; Wang et al., 2013; Zhou et al., 2012; Zhou et al., 2015b; Zhou et al., 2015c). Among the expected and reported LINC and LINC-associated proteins, our endogenous co-IP detected SUN, WIPs, WIT, NCH, and RAN cycle proteins.

In relation to the cytoplasmic and ONM side of the LINC complex, we detected proteins predicted to bind directly to ZmSUN2 including MLKG1 and MLKS2. We also co-precipitated proteins predicted to bind indirectly to ZmSUN2 including MLKT2, two RAN cycle proteins, RAN GTPase, RAN-GAP2, Tub4, Tub6, and the nuclear pore proteins NUP133, NUP96, NUP88, GP210, and NUA. Notably, we did not detect all of the nuclear pore proteins, consistent with previous studies showing interactions of a specific and limited subset of NPC proteins with SUN-domain proteins in association with transcriptional activation and mRNA export processes (Gerber et al., 2004; Li and Noegel, 2015). In relation to the nucleoplasmic and INM side of the LINC complex, we detected two INM proteins, ZmSUN1 and ZmSUN2; and two NMCP/CRWN proteins, NCH1 and NCH2.

The co-IP experiments yielded an interesting list of additional proteins beyond the core LINC and associated proteins summarized in our working model. Some of these proteins appear to belong to non-specific protein categories but in fact have been found in various NE studies involving nuclear actin (ARP7/Actin-1), transcription factors (bZIP5, ARF10, MYBR35, bHLH72/bHLH041), Histone H1 (H1a), and ribosomal proteins (L19, L6, and L22-2) (Horigome et al., 2011; Hotta et al., 2007; Mochizuki et al., 2017; Nakayama et al., 2008). Nuclear actin is increasingly recognized as playing key roles in chromosome dynamics and gene expression (reviewed by Spichal and Fabre, 2017) whereas a plant linker H1 histone was recently shown to have MTOC activity and reside at the nuclear periphery (Boruc et al., 2012; Nakayama et al., 2008).

Given the increasing recognition that the NE and nucleoskeletal proteins are implicated in chromatin remodeling and gene regulation, these additional non-LINC proteins discovered through co-IP represent ideal experimental targets for investigating regulatory mechanisms of chromatin structure and nuclear architecture. For instance, the NE cooperates with nucleoskeletal proteins to modulate euchromatin and heterochromatin organization (Poulet et al., 2017b; Wang et al., 2013). Emerging breakthroughs in HiC mapping of nuclear architecture are likely to uncover additional principles of genome organization that involve LINC-binding nuclear proteins (Grob et al., 2014). Given the importance of lamins in human health, it is possible that plant NMCP/CRWN/NCH defects may shed light on NE-associated diseases.

Both co-IP and FRAP provide complementary assays for investigating *in planta* binding of maize KASH candidates to ZmSUN2 (Fig. 6). For example, MLKG1 was co-purified from maize extracts using SUN2 antisera (Table 2) while FRAP assays demonstrated a ZmSUN2-dependent change in the mobility of GFP-MLKG1 and GFP-MLKP2. The heterologous expression system described in this and previous studies (Zhou et al., 2012; Zhou et al., 2014) represents an efficient means to detect, isolate, and dissect protein-protein interactions for distantly related grass species such as maize, which is not easily transformed. It also provides an efficient means to carry out *in vivo* tests of candidate interactors (e.g. Fig. S6) for which biochemical amounts of native tissue may be limiting.

Overall, this study provides the first comprehensive list of maize LINC candidates (Fig. 1), several of which were validated with multiple lines of evidence. Through the identification of an ensemble of LINC components in a model eukaryote, we have established valuable new avenues of research to address unanswered questions regarding the role of the nuclear envelope as an integrative structure coordinating nuclear and cellular functions.

## METHODS

### Bioinformatic identification of MLKs

The maize B73 proteome v4 (ftp://ftp.gramene.org/pub/gramene/CURRENT_RELEASE/fasta/zea_mays/pep/) was searched for sequences ending in motif [DTVAMPLIFY][VAPIL]PT (brackets indicate alternative amino acid residues at the respective position). The selected proteins were further searched for the presence of TM domain using TMHMM (Krogh et al., 2001) and Phobius software (Käll et al., 2004). Proteins lacking TM domain or specific ending were discarded. The selected maize MLKs were then searched on PLAZA monocot v4 (https://bioinformatics.psb.ugent.be/plaza/versions/plaza_v4_monocots/) and those belonging to same homolog family were grouped together.

### Protein feature calculations and domain searches

Theoretical molecular weight and isoelectric point (pI) of candidate proteins was calculated using the pI/Mw tool of ExPASy (https://web.expasy.org/compute_pi/). Coiled coil domains were identified by Jpred (http://www.compbio.dundee.ac.uk/jpred/). For the identification of conserved domain sequences, the candidate protein sequences were searched on Uniprot (http://www.uniprot.org/), NCBI Conserved Domain Database (https://www.ncbi.nlm.nih.gov/cdd/), PFAM (https://pfam.xfam.org/), and SMART (http://smart.embl-heidelberg.de/). The domains identified were then plotted using IBS 1.0.3 (Illustrator for Biological sequences) software (Liu et al., 2015).

### Identification of LINC orthologs in other organisms and phylogenetic reconstruction

Sequences of orthologs of Maize LINC candidates belonging to matching HOMO gene families were retrieved from PLAZA v4 (https://bioinformatics.psb.ugent.be/plaza/versions/plaza_v4_monocots/). Sequences of all species selected for this study are present in PLAZA except *Medicago truncatula*, which were retrieved from Gramene. The protein sequences were further screened for the presence of characteristic protein features. The MLKS/MLKG/MLKP orthologs were searched for the presence of TM domains towards C-terminus and for terminal four residues of the proteins to be VIPT, VVPT, AVPT, PLPT, TVPT, LVPT or PPPS. For MLKT orthologs, proteins having high sequence similarity and C-terminal TM domains were retained. For NCHs, KAKUs and NEAPs, sequences having high sequence similarity to Arabidopsis CRWNS, KAKU4 and NEAPs were selected. For phylogenetic reconstruction of each protein category, sequences were first aligned using MUSCLE and Neighbour end joining trees were generated in MEGA 7.0.21 (Kumar et al., 2016) using default parameters.

### Transcriptome

The expanded maize gene expression atlas (Stelpflug et al., 2016) was downloaded for analysis. Log10 Fragments Per Kilobase of exon per Million fragments mapped (FPKM) values for 15 tissues were used to construct a heatmap using the Heatmap2 package of R. To group tissues by similar expression profiles, hierarchical clustering was done using the Hclust function of R.

### Co-IP protocol

Maize (Zea mays L) B73 seeds were obtained from WF Thompson (NC State University, Raleigh, NC, USA, accession “27 FARMS”, bulked for NCSU, denoted B73NC27). Seeds were grown at the Florida State University Mission Road Research Facility (Tallahassee, FL). Young earshoots (3-5 cm long) were harvested from 7-8-week-old plants, flash frozen in liquid nitrogen and stored at −80 °C until use. Ten grams tissue was ground under liquid nitrogen with a mortar and pestle, and cross-linked with 0.1% formaldehyde by stirring for 10 min in 100 mL ice-cold Buffer A (15 mM PIPES-NaOH, pH 6.8, 0.32 mM sorbitol, 80 mM KCl, 20 mM NaCl, 0.5 mM EGTA, 2 mM EDTA, 1 mM DTT, 0.15 mM spermine, and 0.5 mM spermidine; (Belmont et al., 1987). The nuclei were isolated as described in (Vera et al., 2014) and resuspended in 5 ml PBS. The isolated nuclei were treated with 2 mM DSP (dithiobis(succinimidyl propionate) (Cat. No. 22585, Thermo Scientific) crosslinker for 2 h at 4 °C. To lyse the nuclei, 7 M urea, and 1% Triton X-114 and plant protease inhibitor cocktail (Cat. No. P9599, Sigma) were added and incubated for 20 min on ice. The ruptured nuclei were sonicated three times in Bioruptor (Diagenode) for 5 minutes each at medium setting. Urea was diluted with 8 volumes of PBS. Homogenates were centrifuged at 10,000 xg for 10 min at 0 °C to remove cellular debris. The supernatant was collected and pre-cleared by adding 0.5 mg Protein A dynabeads (Cat No.10000D, Thermo Fisher) for 30 min with rotation at 4 °C. The beads were removed by placing tubes on magnetic rack (Cat. No.12321D, Thermo Fisher). The cleared lysate was split into 3 equal aliquots, and 3 μg α-SUN2 antibody was added to first aliquot, 3 μg NDPK control antibody to the second tube and no antibody added to the third tube. The tubes were incubated at 4 °C with rotation overnight, then 0.5 mg Protein A magnetic beads were added to each tube for one-hour incubation at 4 °C. The antigen-antibody bound beads were captured with the help of a magnetic rack and washed thrice with PBS supplemented with 0.02% Tween-20. The protein complexes bound to the beads were digested using ProteoExtract All-in-One Trypsin Digestion Kit (Cat. No. 650212, Calbiochem) according to manufacturer’s instructions. Briefly, beads were resuspended in extraction buffer and digestion buffer was added, then reduced using the provided reducing agent for 10 minutes at 37°C. Samples were cooled to RT and then blocked using blocking reagent for 10 minutes at room temperature. Trypsin at a final concentration of 8 ng/μl was added and incubated for 2 hours at 37°C with shaking. The reaction was stopped by acidification. Digests were vacuum dried and submitted for LC-MS/MS to the Translational Lab at Florida State University.

### Mass spectrometry and data analysis

Digested peptides were resuspended in 0.1% Formic acid for loading onto LC-MS at the Translational Lab, Florida State University. An externally calibrated Thermo LTQ Orbitrap Velos nLC-ESI-LTQ-Orbitrap (high-resolution electrospray tandem mass spectrometer) was used with the following parameters. A 2 cm, 100 μm i.d. (internal diameter) trap column (SC001 Easy Column from Thermo-scientific) was followed by a 10 cm analytical column of 75 μm i.d. (SC200 Easy Column from Thermo-scientific). Both trap column and analytical column had C18-AQ packaging. Separation was carried out using Easy nanoLC II (Thermo-Scientific) with a continuous, vented column configuration. A 5 μL (~500 ng) sample was aspirated into a 20 μL sample loop and loaded onto the trap. The flow rate was set to 300 nL/min for separation on the analytical column. Mobile phase A was composed of 99.9 H2O (EMD Omni Solvent) and 0.1% formic acid, and mobile phase B was composed of 99.9% ACN and 0.1% formic acid. A 1 h linear gradient from 0% to 45% B was performed. The LC eluent was directly nano-sprayed into an LTQ Orbitrap Velos mass spectrometer (Thermo Scientific). During the chromatographic separation, the LTQ Orbitrap Velos was operated in a data-dependent mode and under direct control of the Xcalibur software (Thermo Scientific). The MS data were acquired using the following parameters: 10 data-dependent collisional-induced-dissociation (CID) MS/MS scans per full scan. All measurements were performed at room temperature and three technical replicates. The raw files were analyzed using Proteome Discoverer (version 1.4) software package with SequestHT and Mascot search nodes using the maize proteome database available at Gramene (AGVP3.0). The resulting msf files were further analyzed by the ‘Scaffold version 4.4’ proteome validator software at 90% peptide and protein threshold with a minimum of 1 peptide.

### Constructs and gene cloning

Coding sequences of MLKP1 and MLKP2 genes fused to the C-terminus of eGFP-FLAG-HA sequence were custom synthesized by Genscript as pHWBF08 and pHWBF06 respectively. Sequences eGFP-FLAG-HA-MLKP1 and eGFP-FLAG-HA-MLKP2 were PCR amplified with att flanking primers and cloned individually into pDONR221 vector by BP cloning (Cat. No. 1235019, Invitrogen) to generate pHWBF08EC and pHWBF06EC entry clones respectively. These were moved to destination vector pH7WG2 (Karimi et al., 2002) by LR recombination reaction of Gateway cloning system (Cat. No. 1235019, Invitrogen) to obtain expression vectors pPK1iexp and pPK1Cexp respectively. For MLKG1, the coding region was PCR amplified from maize B73 using gene specific primers (Table S4) and cloned into pCRII vector by TOPO cloning (Cat. No. K461020, Thermo Fisher). After sequence verification, the coding sequence was amplified with forward primer flanked with BamHI restriction site sequence and reverse primer flanked by SbfI restriction site sequence. The PCR amplified product was double digested with BamHI and SbfI (NEB) and ligated in pECGFP (Bass lab) donor vector in frame and downstream of the eGFP-FLAG-HA sequence. The fusion sequence was then moved to pH7WG2 destination vector by LR recombination to generate an expression vector called pPK1Dexp. Deletion constructs for MLKP1 and MLKP2 were generated using corresponding full length entry clones as template to PCR amplify required regions with internal primers containing flanking att sequences (Table S4) with stop codon and cloned into pDONR211 by BP cloning. These deletion constructs were subcloned in pH7WG2 destination vector by LR recombination. MLKG1ΔKASH was generated using the Q5 DNA mutagenesis kit (Cat. No. E0554S, NEB). Briefly, two primers were designed in opposite directions, going outwards from the KASH domain. The entire entry vector (pECPK1D) minus the KASH domain was amplified using high fidelity Q5 polymerase provided with the kit. The template contamination was removed and vector re-ligated using manufacturer-provided enzyme mix containing a kinase, a ligase and DpnI to create entry clone MLKP1delKASH. The entry clone was then moved to pH7WG2 destination vector by LR recombination. Sequences of all the plasmids generated in this study are given in Table S5.

### *Agrobacterium tumefaciens* transformation

Using established heat shock protocols, *A. tumefaciens* was transformed with expression vectors (Graumann and Evans, 2010). Briefly, competent cells were thawed on ice and 4ul plasmid DNA was added. Cells were incubated for 5 min on ice, 5 min in liquid nitrogen, and 5 min at 37 °C, then 1 ml LB medium was added and cells were incubated for 2h at 28 °C before being plated out and left to grow for 48 h at 28 °C.

### *Nicotiana benthamiana* transformation and microscopy

Transient expression of fluorescent fusion proteins was carried out as previously described (Sparkes et al., 2006). Briefly, Agrobacterium were suspended in infiltration buffer at an OD of 0.05 and pressed into the lower leaf epidermis with a syringe. Three days post infiltration, leaf sections were imaged with a Zeiss LSM 800 confocal microscope using a 100X oil immersion objective, NA 1.46 and 1.5X digital zoom. GFP was excited at 488 nm and emission collected between 500-550 nm; mCherry was excited with a 561 nm laser and emission collected from 565-620 nm. For FRAP, we performed 5 pre scans and then a 160×160 pixel ROI was bleached with the 488 nm laser at 100% for 10 iterations. Recovery data was collected over 36 s (240 scans, 0.13 s framerate). Mean ROI intensity data was extracted using Zen Blue (version 2.3) and normalised as described in (Martinière et al., 2012). One phase association recovery curves, plateau values and halftimes were produced in Graphpad (version 6). The ANOVA statistical test was performed with multiple comparisons. For DAPI staining, leaf sections were first incubated in 1 ml water containing 5 ul 20% Triton X-100 and 1ul DAPI. DAPI excitation was performed with the 405 nm laser and emission between 400-450 nm imaged. DAPI/GFP or DAPI/mCherry imaging was performed using multitrack line scanning.

### Direct Co-IP and western blot

Leaves of *N. benthamiana* (1 g) were harvested and ground in liquid nitrogen. The powdered tissue was homogenized in 5 mL of radioimmunoprecipitation assay buffer (50 mM Tris-HCl, pH 7.5, 150 mM NaCl, 0.1% SDS, 0.5% Na-deoxycholate, 1% NP-40, 1 mM PMSF, and 1% protease inhibitor cocktail (Cat. No. P9599, Sigma) in a dounce homogenizer kept on ice. The homogenate was centrifuged at 10,000 xg and the supernatant was collected for co-IP. One tenth of the protein extract was used as the input sample and concentrated using acetone precipitation. The rest of the lysate was subjected to co-IP using dynabeads coated with rabbit anti-GFP antibody (Cat. No. ab290, Abcam). The lysate was incubated with beads for 4 hours, after which the beads were collected using magnetic rack and washed with PBST three times. The antibody-protein complex bound to beads was eluted using 50 mM Glycine (pH 2.8). The immunoprecipitates and the input samples were boiled in 4X Laemelli’s gel loading buffer, fractionated on 15% SDS-PAGE and transferred to polyvinylidene difluoride membranes (Cat. No. 88518. Thermo Scientific), and detected with a mouse anti-GFP (1:3,000; Cat No. MA5-15256,Thermo Scientific) or rabbit anti-SUN2-NPAP (1:1,000; Murphy et al 2014) antibody at room temperature for 1 h. After three 15-min washes in PBS-T buffer at room temperature, the membranes were incubated with a 1:3,000 dilution of anti-rabbit or anti-mouse IgG horseradish peroxidase–linked antibody (Santa Cruz Biotechnology, Santa Cruz, CA) for 1 h at room temperature, then subjected to 3 × 15 min washes in PBST buffer at room temperature. The PVDF membrane was incubated with chemiluminescent reaction kit for 3-5 min at room temperature (Millipore, Immobilon detection kit, WBKL50100, Billerica, MA) and exposed to X-ray film in dark room to visualize the protein bands.

## ACKNOWLEDGEMENTS

Thank Eric Richards and Iris Meier for helpful and insightful comments on the manuscript, Allison Jevitt for artwork on Figure 1, Dr. Rakesh K Singh and staff of the Translational Lab (College of Medicine, FSU) for assistance and advice on co-IP and Mass spectrometry. The work was supported by grants from the National Science Foundation to HWB (NSF IOS 1444532) and Oxford Brookes University to KG, and from a Dissertation Research Grant to HKG (Florida State University Graduate School).

## Supplementary Figure legends

**Figure S1.**
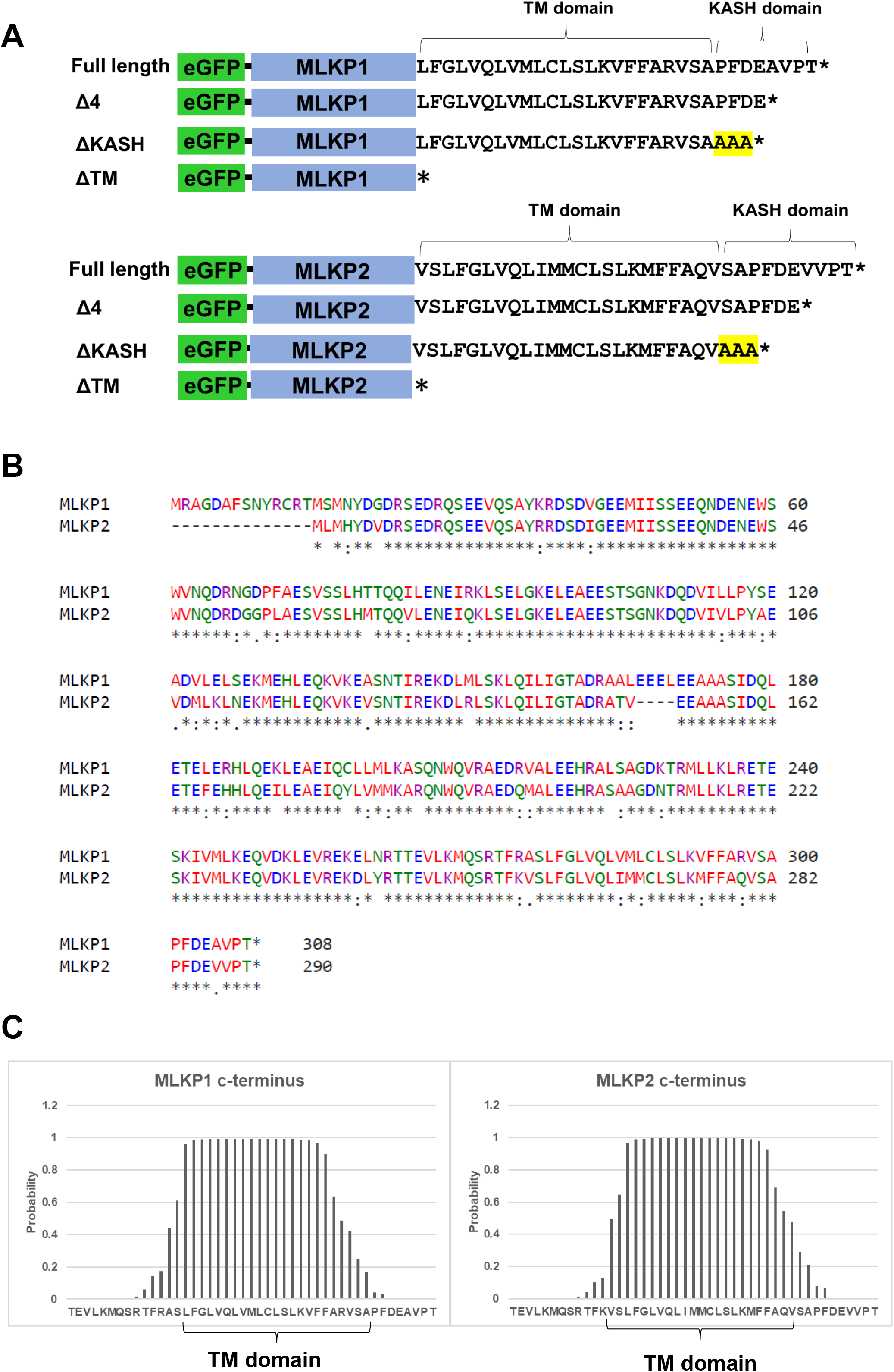
MLKP1 and MLKP2 deletion constructs. The fluorescent fusion protein (eGFP) constructs for MLKP1 and MKLP2 are depicted. (A) Schematic representation showing the location of the transmembrane and KASH domains in full length and deletion constructs. Three alanine residues added to the C-terminus of ΔKASH constructs are highlighted in yellow. (B) Pairwise protein sequence alignment (Clustal Omega) of MLKP1 and MLKP2 is shown. (C) Posterior probabilities (Phobius transmembrane topology predictor) of TM helix for C-termini of MLKP1 and MLKP2 are plotted.

**Figure S2.**
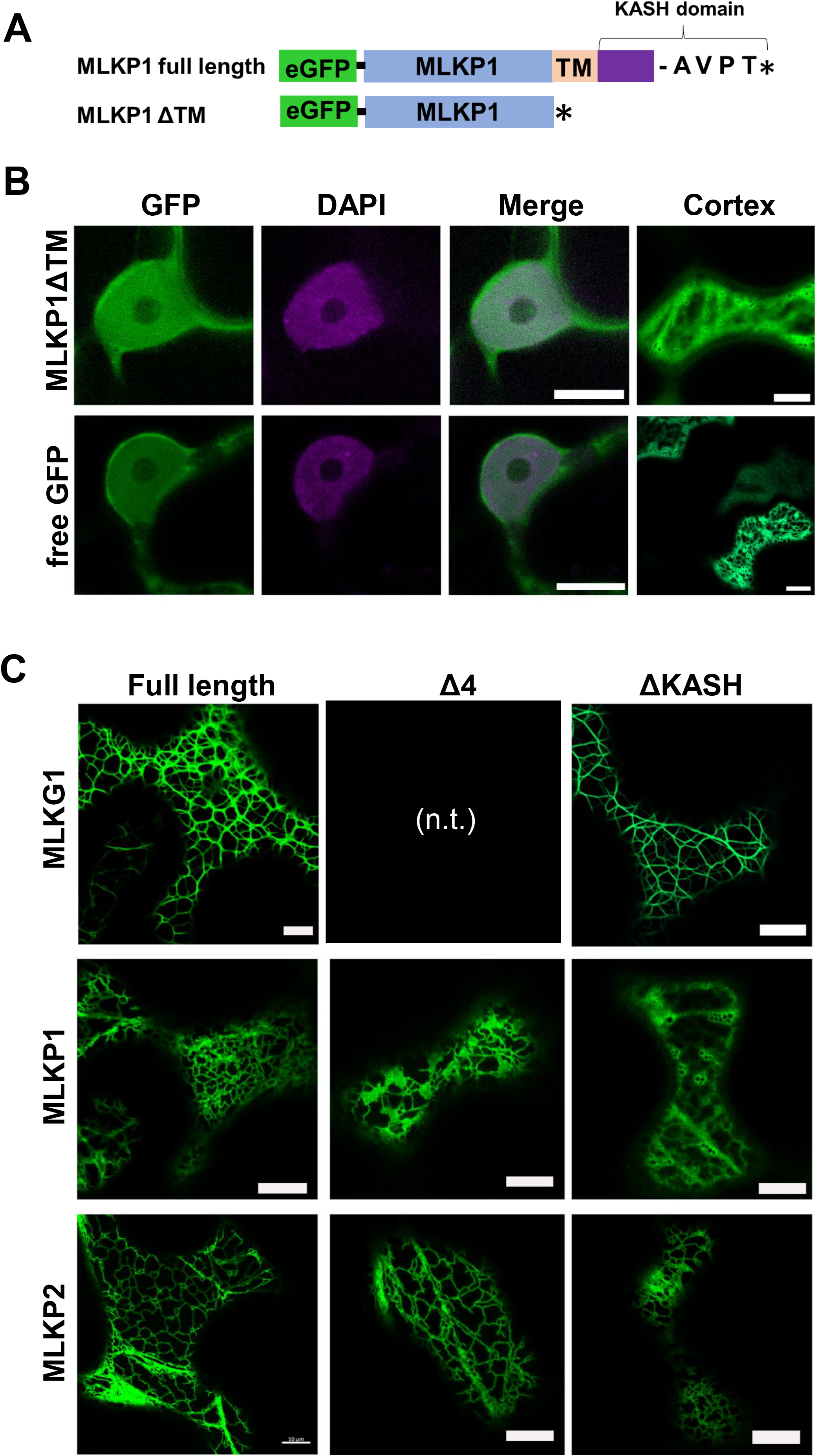
Additional subcellular localization staining patterns of GFP-KASH constructs from *N. benthamiana* transient expression microscopy. GFP localization assays were performed as described for Figure 5. (A) Schematic overview of domain composition of the MLKP1 constructs. (B) Staining patterns in and around the nucleus (1st three columns) and a the cell periphery (last column) shows that the construct lacking the TM domain and the GFP only are distributed throughout the cytoplasm and nucleoplasm. Scale bars for panel B are 5 μm. (C) Cell periphery images show the cytoplasm near the cell cortex for full length and domain deletion constructs. Gene names are listed on the left for each row; constructs are listed on the top for each type; n.t. = not tested). The MLKP1 terminal deletions (both Δ4 and ΔKASH) result in a more soluble staining pattern compared to the full length. The other deletions retain a pattern typical of ER association. Scale bars are 10 μm.

**Figure S3.**
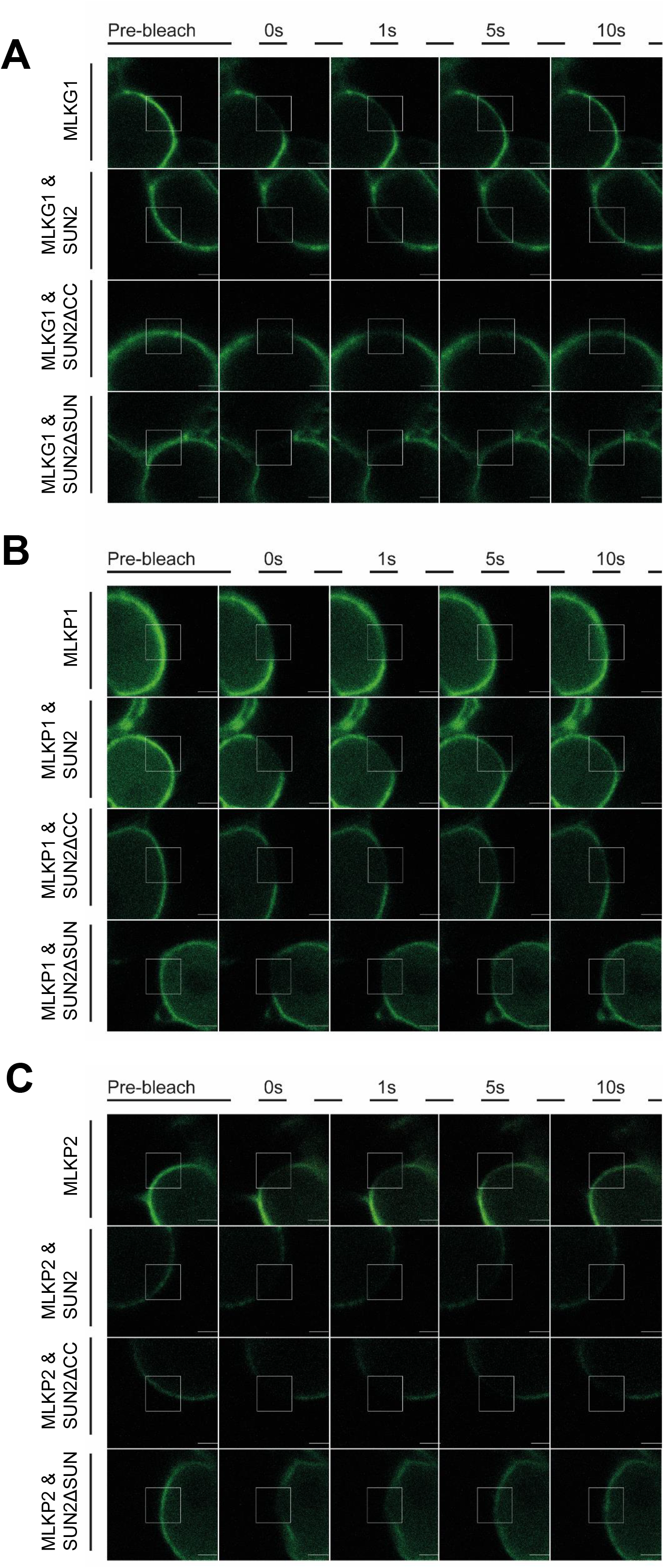
FRAP images from the co-expression interaction assays. Representative images of cells during the FRAP assays for the experiments summarized in Figure 6. The FRAP ROIs (white boxes) are shown along with images taken at the times indicated across the top for (A) GFP-MLKG1, (B) GFP-MLKP1, and (C) GFP-MLKP2. Scale bar denotes 2μm. FRAP ROI shown by white box. All scale bars 2μm.

**Figure S4.**
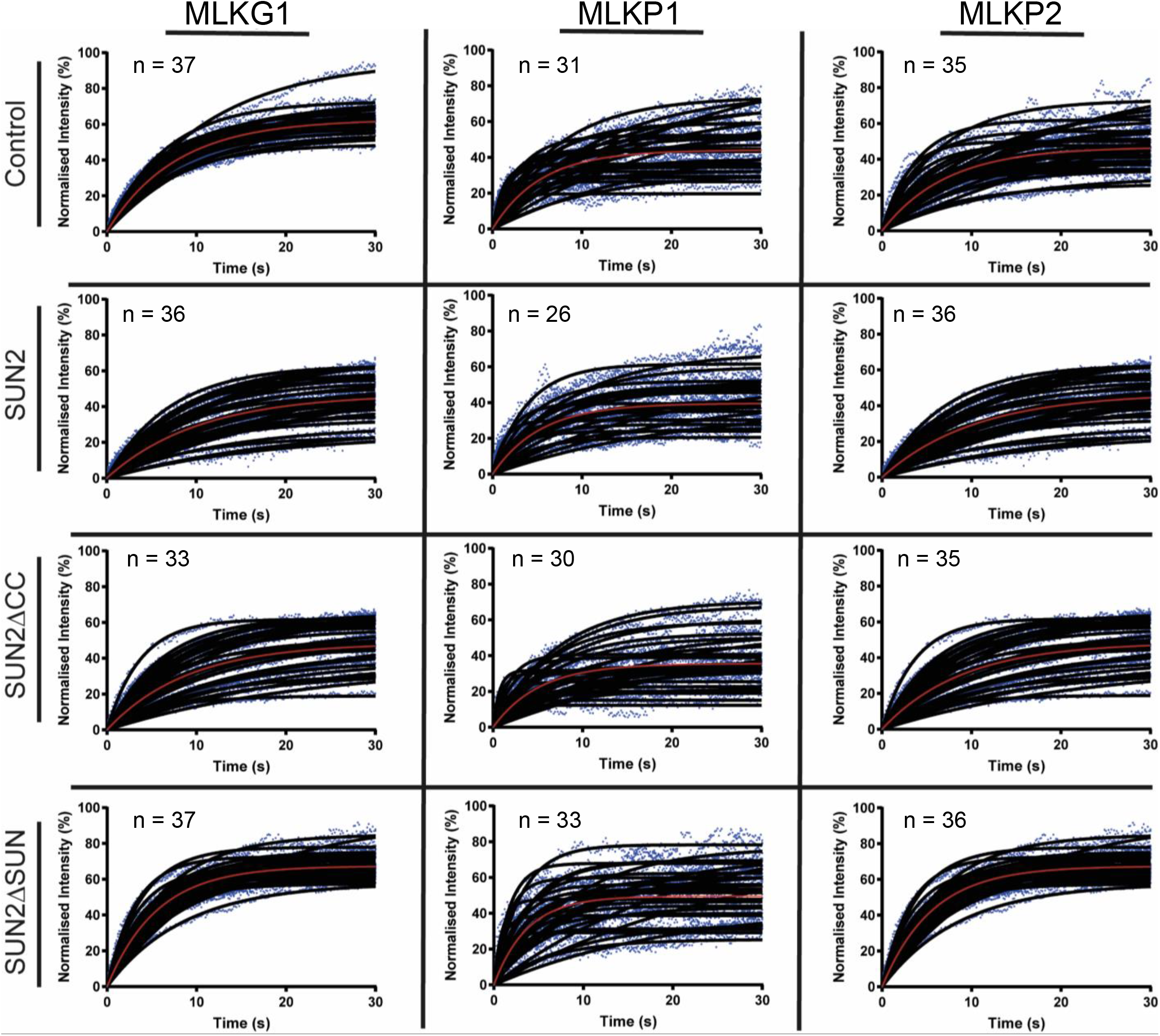
Normalized intensity FRAP curves co-plotted from all experiments performed in *N. benthamiana* leaves. The GFP fusion protein constructs (MLKG1, MKLP1, and MLKP2) are grouped by columns as labeled at top. The co-expression constructs are grouped by row as labeled at left. Individual data points (blue dots), single experiment recovery curves (black lines) and global average recovery curves (red lines) and replicate numbers (n) are shown in each panel.

**Figure S5.**
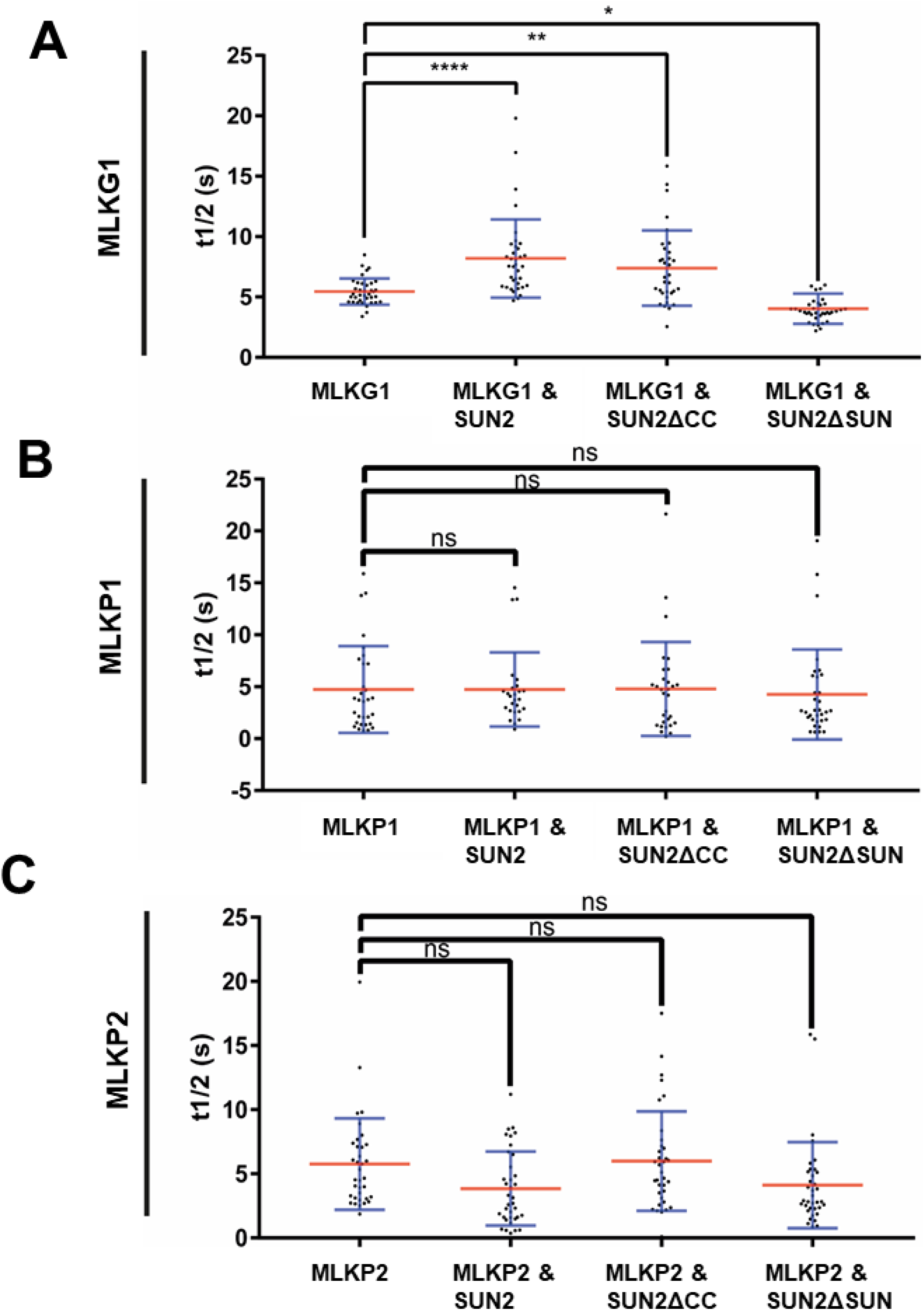
FRAP recovery halftime values. The recovery half times (t1/2(s)) are plotted for the full-length and co-expression experiments summarized in Figure 6. The SD (blue lines) and mean (red lines) are shown for (A) GFP-MLKG1, (B) GFP-MLKP1, and (C) GFP-MLKP2. ANOVA statistical analyses with multiple comparisons are noted (ns = not significant = p≥0.05, ** = significant with P ≤0.01, **** = significant with P ≤0.0001). Replicate numbers range from n=26 to n=35, as listed in Supplemental Figure 3.

**Figure S6.**
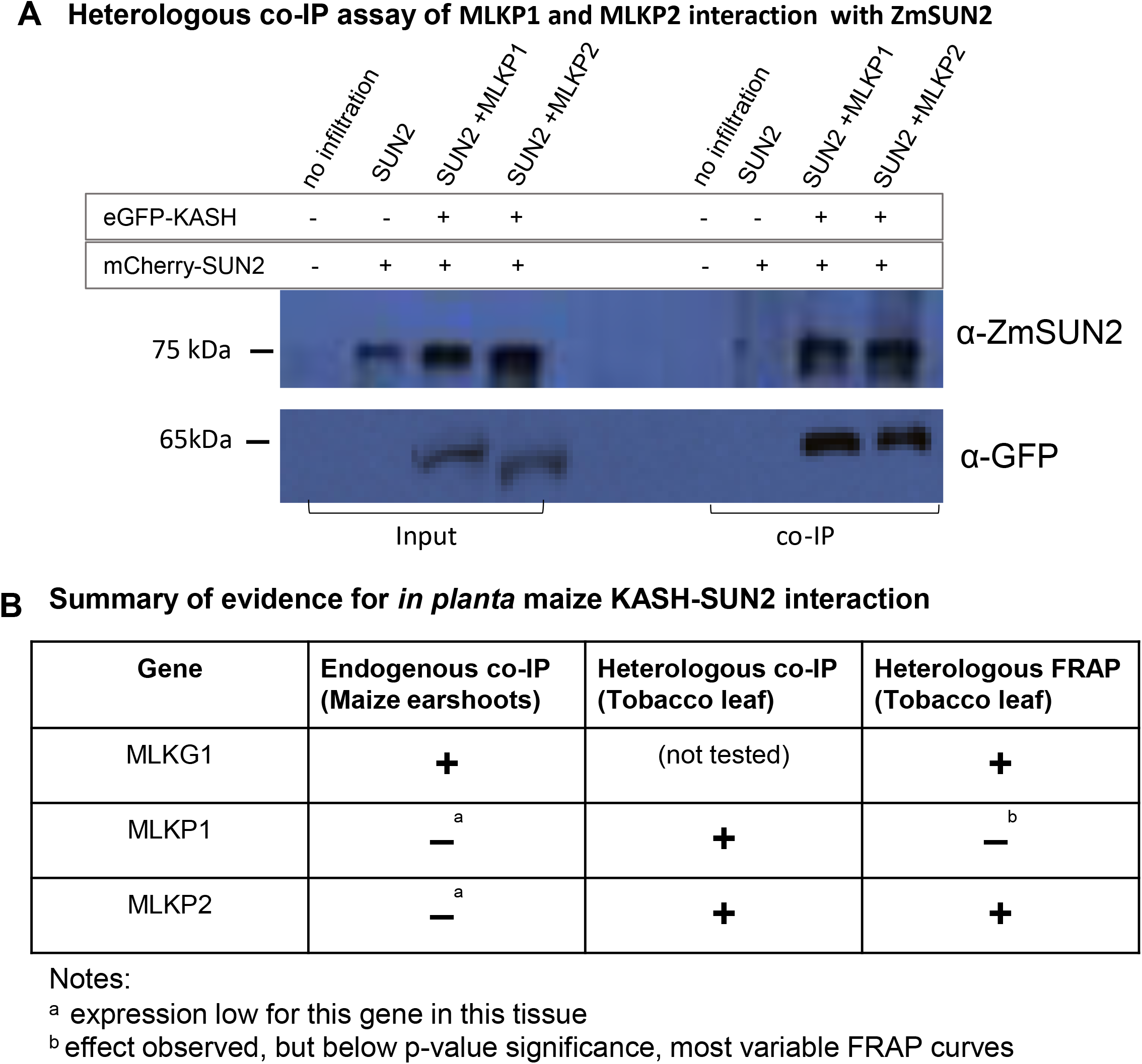
Co-immunoprecipitation assays of MLKP1 and MLKP2 with ZmSUN2 in *N. benthamiana*. Total protein extracts from *N. benthamiana* leaves harvested 3 days post-infection were subjected to co-IP using dynabeads coated with anti-GFP antibody. (A) Western Blot of the co-precipitates and inputs with primary antibodies listed on the right side of the blot. The FP construct combinations used for infiltration are listed on the top of the blot. The numbers on the left are protein molecular weight standards. (B) A summary of KASH-SUN interaction evidence tabulated from this experiment (tobacco heterologous co-IP) or other experiments in this study (maize endogenous co-IP from Table 2, Heterologous FRAP form Figure 6 and Supplemental Figures 2-4).

**Figure S7.**
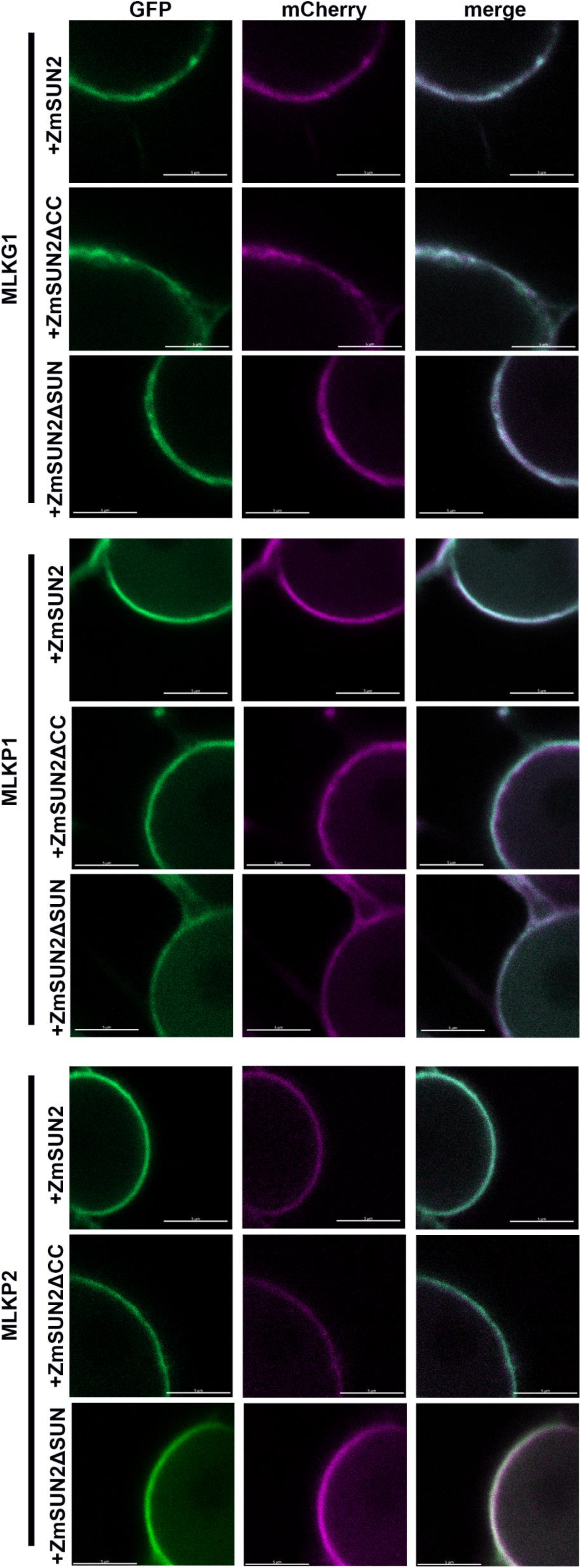
Co-localization of proteins from GFP-KASH and mCherry-SUN constructs. Co-localization of GFP-MLKG1, GFP-MLKP1 and GFP-MLKP2 (green) with mCherry-ZmSUN2, mCherry-ZmSUN2ΔCC and mCherry-ZmSUN2ΔSUN (magenta) at the nuclear periphery. Size bar 5μm.

## List of Symbols and Abbreviations

CDS: coding sequence
CRWN1: CROWDED NUCLEI 1s
FRAP: Fluorescence Recovery After Photobleaching
INM: inner nuclear membrane
KASH: Klarsicht/ANC-1/Syne-1 homology
LINC: linker of nucleoskeleton and cytoskeleton
MLKP: Maize LINC KASH AtWIP-like
MLKG: Maize LINC KASH Grass-specific
MLKS: Maize LINC KASH AtSINE-like
MLKT: Maize LINC KASH AtWIT-like
MKAKU4: Maize KAKU4-like
NE: nuclear envelope
NET: nuclear envelope transmembrane protein
NCH: NMCP/CRWN-Homologous
NEAP: nuclear envelope-associated protein
ONM: outer nuclear membrane
SUN: Sad1/UNC-84
TM: transmembrane
WIP: WPP domain-interacting protein
WIT: WPP domain-interacting tail-anchored protein

